# *Whippet*: an efficient method for the detection and quantification of alternative splicing reveals extensive transcriptomic complexity

**DOI:** 10.1101/158519

**Authors:** Timothy Sterne-Weiler, Robert J. Weatheritt, Andrew Best, Kevin C. H. Ha, Benjamin J. Blencowe

## Abstract

Alternative splicing (AS) is a widespread process underlying the generation of transcriptomic and proteomic diversity in metazoans. Major challenges in comprehensively detecting and quantifying patterns of AS are that RNA-seq datasets are expanding near exponentially, while existing analysis tools are computationally inefficient and ineffective at handling complex splicing patterns. Here, we describe *Whippet*, a method that rapidly, and with minimal hardware requirements, models and quantifies splicing events of any complexity without significant loss of accuracy. Using an entropic measure of splicing complexity, *Whippet* reveals that approximately 33% of human protein coding genes contain complex AS events that result in substantial expression of multiple splice isoforms. These events frequently affect tandem arrays of folded protein domains. Remarkably, high-entropy AS events are more prevalent in tumour relative to matched normal tissues, and these differences correlate with increased expression of proto-oncogenic splicing factors. *Whippet* thus affords the rapid and accurate analysis of AS events of any complexity, and as such will facilitate biomedical research.

## Introduction

High-throughput RNA sequencing (RNA-seq) technologies are producing vast repositories of transcriptome profiling data at an ever expanding pace^1^. This explosion in data has enabled genome-wide investigations of the role of alternative pre-mRNA splicing (AS) in gene regulation and dysregulation. Initial investigations of tissue transcriptomes using RNA-seq data revealed that more than 95% of human multiexon genes undergo AS^2,3^. These and more recent studies analyzing ribosome-engaged splice-variant transcripts suggest that AS is potentially the most widespread process underlying the generation of transcriptomic and proteomic complexity^4–6^. Furthermore, numerous AS events belonging to co-regulated exon networks have been shown to provide critical functions in diverse normal and disease-associated processes and pathways in multicellular organisms^7^.

A major challenge confronting the advancement of knowledge of AS-complexity regulation and function is that existing methods for analyzing RNA-seq data require extensive computational resources and expertise. The first steps in the analysis of AS using RNA-seq data involve the alignment of reads to a transcriptome or reference genome, followed by quantification of AS levels by downstream methods. Both steps can be time-consuming. For example, aligning 50 million paired-end reads with the widely-used program Tophat2^8^ takes over 15 hours on 15 processing cores, and quantification of AS using the aligned reads can take an additional seven hours using Mixture-of-Isoforms (MISO), one of the most highly cited AS analysis tools^9^. Because RNA-seq data is being generated at a rate that is outpacing parallel advancements in computational hardware required for analysis, new methods that are both efficient and accurate are necessary.

To address these challenges, recent developments in transcript-level expression quantification have circumvented the read alignment step by extracting and quantifying k-mers (i.e. all possible subsequences of length k) from reads, which can decrease processing times by over 20 fold^10,11^. Unfortunately, transcript-level quantification is not well-suited for the analysis of splice isoforms. This is because transcript-level quantification assumes dependency between multiple alternatively processed sequences of a gene. However, these regions are often regulated independently such that the assumption of dependency is invalid. (Supplemental Fig. 1a). Short read-based RNA-seq analyses therefore require that event-level approaches are used for the quantification of AS. While considerable advancements have been made on this front^12^, most existing tools analyze individual AS events using simple binary models (i.e. a single alternative exon surrounded by two constitutive exons) and are not suited for the analysis of relatively complex AS patterns.

Accordingly, an important goal for understanding how transcriptomes shape key biological processes and pathways is to develop new analysis methods that are capable of efficient detection and quantification of both simple and complex splicing patterns. To address these challenges, we have developed *Whippet*, an easy to use software for the rapid detection and quantification of AS events of any complexity that has computational requirements compatible with a laptop computer. We demonstrate the utility of Whippet in the discovery of previously unknown AS complexity in vertebrate transcriptomes associated with the regulation of tandem domains and other protein sequence features, as well as an unexpected increase in AS complexity and entropy in cancer transcriptomes.

## Results

### Using contiguous splice graphs to quantify alternative splicing at the event-level

To facilitate event-level quantification, Whippet creates ‘*Contiguous Splice Graphs*’ (CSGs) using transcript features extracted from gene annotations. These are directed graphs whose nodes are exonic sequences and whose edges can either be spliced (intron excision) or unspliced (between two adjacent nodes). Single isoforms transcribed from a gene represent individual paths through the CSG. Representing transcript architecture in this manner thus allows a vast number of different isoforms from a gene to be represented by a single CSG.

Whippet aligns reads to exon-exon junction-spanning sequences represented by a CSG to detect pre-mRNA splicing events. For each CSG, adjacent non-redundant exonic sequences (nodes) are arranged into a single sequence (Fig. 1a, Supplemental Fig. 1b-d). The CSG sequences are concatenated and a global transcriptome Full-text index in Minute space (FM-index)^13^ is built, which compresses the input data and facilitates efficient full-text pattern searching (Fig. 1a). Raw sequence reads are seeded to the transcriptome FM-Index, and alignments are extended in the forward and reverse directions (Fig. 1b). Nucleotide k-mers flanking each annotated 5’ or 3’ splice-site are used as an index for two hash-tables (i.e. associative maps) that link to gene-node pairs. Reads intersecting two sets of (gene, node) pairs for an exon-exon junction sequence (upstream 9mer + downstream 9mer = exon-exon junction 18mer by default) produce compatible nodes that are recursively extended along the CSG for the remaining length of the read (Fig. 1b, Supplemental Fig. 2; see Methods). Performing read alignments in this manner affords Whippet considerable efficiency by storing minimal data while still supporting pseudo-*de novo* AS event identification (see below).

**Figure 1.**
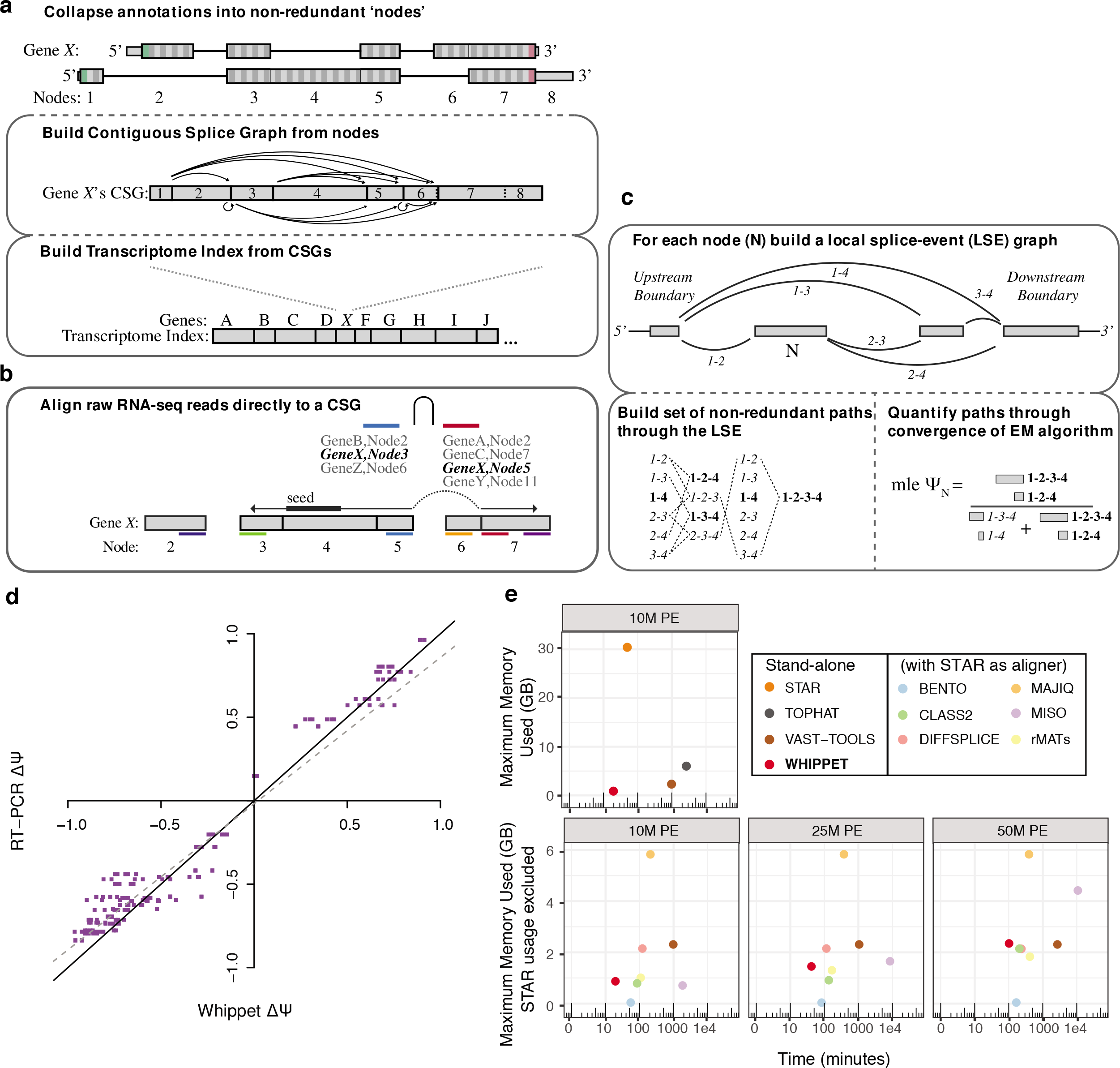
Overview of Whippet algorithm, validation and benchmarking (a) Example gene model with two alternative isoforms and Whippet’s node assignments for the gene (*top*). The Contiguous Splice Graph (CSG) for the same exemplar gene model is shown, with spliced edges as solid lines with arrows, straight solid lines indicating spliced node boundaries, and straight dotted lines indicating unspliced edges between nodes (*middle*). A single transcriptome Full-text index in Minute space (FM-Index) is built from concatenated CSG sequences, with solid lines indicating separation between each CSG (*bottom*). **(b)**Diagram of the Contiguous Splice Graph Alignment (CSGA) algorithm. An alignment is seeded from a raw RNA-seq read, and then extended in both directions. Spliced extension utilizes a k-mer-indexed associative map of (gene, node) pairs and sorted intersection to efficiently return the set of compatible spliced edges (see Methods). **(c)**Example of a Whippet analysis of a local splicing event for node N, defined as the set of edges between the two boundary nodes (*top*). To determine the Percent Spliced In (Ψ) of some node N, the full set of non-redundant paths through the local splice event are enumerated (*bottom left*), and then quantified through convergence of the expectation-maximization (EM) algorithm (*bottom right*) (see Methods). mle, maximum likelihood estimation **(d)**Comparison of ΔΨ (change in PSI) values between human liver and cerebellum as derived from Whippet Ψ predictions from RNA-seq data, and Ψ values from RT-PCR data from the same tissue types. Best fit linear regression line (*dotted line*) and diagonal y=x (*solid line*) are shown **(e)**Comparison of the maximum memory-usage (*y-axis*) and log-scaled time (*x-axis*) requirements of Whippet relative to several published methods for RNA-seq read alignment and splicing quantification. The top panel compares resources used by Whippet with the commonly used RNA-seq aligners TOPHAT^8^ and STAR^14^, and with VAST-TOOLS^15^, when aligning 10 million (M) paired-end (PE) RNA-seq reads. The bottom panels compare resources used by Whippet and six other splicing quantification methods when analyzing aligned reads (10M, 25M or 50M) (see Methods). With the exception of Whippet and VAST-TOOLS, all algorithms in the bottom panel require initial alignment of reads by an aligner from the top panel. GB,gigabyte.

After alignment of all sequencing reads to the CSG transcriptome, Whippet builds each node’s local splicing event (LSE), which is defined as the set of nodes and observed edges between the farthest overlapping or connecting upstream and downstream boundary nodes (Fig. 1c, see Methods). Since observed CSG paths in an LSE may share common nodes or edges, the proportional abundance of each path in the LSE is determined through maximum likelihood estimation using the EM (expectation-maximization) algorithm (see Methods). The percent-spliced-in (PSI, Y) of a node is then defined as the sum of the proportional abundance of the paths containing the node (Fig. 1c).

### Whippet facilitates rapid and accurate analysis of RNA-seq data for low- and high-complexity splicing events

To assess the performance of Whippet, we compared its efficiency and accuracy with established and newer analysis methods, using both simulated and experimental RNA-seq datasets. Whippet’s CSG alignment algorithm correctly maps 94-96% of simulated spliced junction reads, which is within ±2-4% of the percentage of reads correctly mapped by widely used spliced-read aligners such as STAR^14^ and TOPHAT2^8^, although it achieves this task at a faster rate and with lower memory usage than these aligners (Fig. 1e, Supplemental Fig. 3a). To assess the accuracy of AS-quantification, we compared published RT-PCR-derived Ψ values for AS events in liver and cerebellum samples with Whippet-derived Ψ values obtained from RNA-seq data from the same tissue types^12^. Whippet-derived Ψ values correlated highly with the RT-PCR measurements (*r*=0.963, Pearson’s correlation coefficient) and with a comparable error rate as the best other methods tested (Fig. 1d, Supplemental Fig. 4)

We next assessed whether the unique architecture of Whippet increased its speed and performance relative to other AS quantification methods that have a comparable degree of accuracy (Supplemental Fig. 4b, p > 0.05, Wilcoxon rank-sum one sided test, Whippet vs other algorithms). To this end, we compared its speed and memory usage to those of eight other highly-cited splicing tools^9,12,15–19^ when analyzing several paired-end RNA-seq datasets from HeLa cells with increasing read depth (∼10M, ∼25M and ∼50M). Remarkably, Whippet quantifies AS on an event-specific basis from a raw paired-end 25M RNA-seq read dataset in less than forty-five minutes on a typical cluster node (Dual-Core AMD Opteron(tm) Processor 8218, 2.5 GHz, 60GB RAM, 1,024KB cache), and uses less than 1.5GB of memory. This represents a considerable increase in performance over all other event-level tools tested both in terms of speed and memory usage (Fig. 1e, Supplemental Fig. 3a and d). For example, MISO, the most highly-cited splicing tool, took more than 24 hours and used 30 GB of memory to analyze the same data (Fig. 1e). Moreover, on a personal laptop with a solid-state hard drive (Macbook Pro 3.1 GHz Intel i7), Whippet was able to quantify the 25M read paired-end dataset in only 13.4 minutes, whereas this was not feasible with other tools using the same hardware due to excessive RAM usage and computational time. Together, these results demonstrate that Whippet is both a highly efficient and accurate method for the analysis of AS using RNA-seq data.

The benchmarks described so far focus on “simple” AS events, such as single cassette alternative exons that are flanked by two pre-defined constitutive exons and that have binary splicing outcomes. However, some AS events are more complex and can involve situations where a splice site is variably paired with two or more other sites^20^. Whippet’s graph-based algorithm is designed to detect and quantify such AS event complexity in two related ways: first, it classifies events into discrete categories of splicing complexity based on total numbers of possible combinatorial outcomes (i.e. K(*n*) = 2^*n*^ spliced outcomes for K1,…, K6; Fig. 2a, Supplemental Fig. 5), and second, it calculates a Ψ-dependent measure of AS complexity using Shannon’s entropy (i.e., entropy = −Σ_*i*_ Ψ_*i*_ *log*_2_Ψ_i_ such that the maximum entropy for an event in K(*n*) is *n*; Fig. 2b, Supplemental Fig. 6a). This entropic measure of alternative splicing formalizes the total number of possible outcomes for an AS event and the degree of their proportional contribution to the transcriptome in a read-depth and read-length-independent manner (Supplemental Fig. 6b-c)

**Figure 2.**
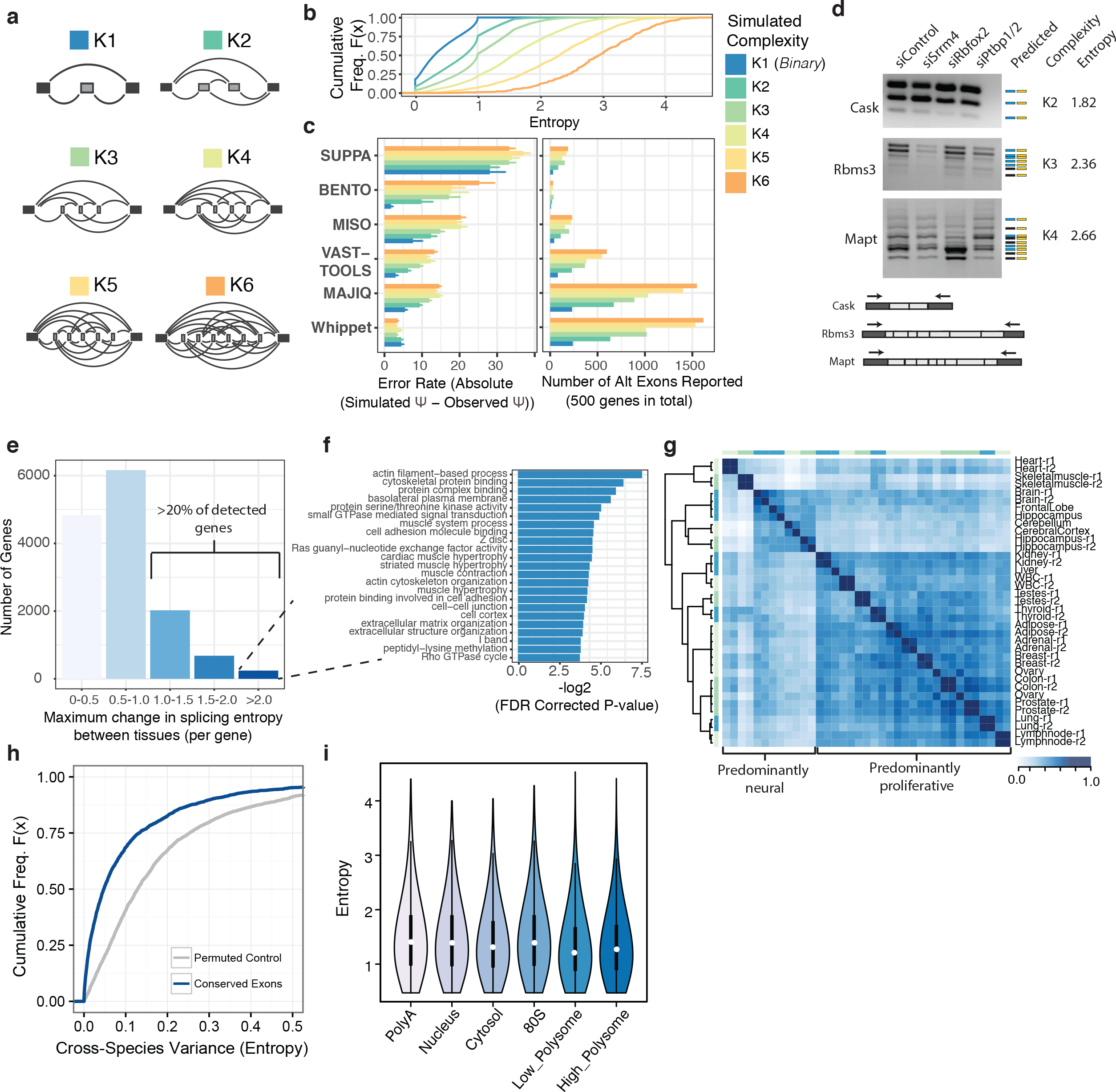
Identification and characterization of high entropy events (a) Formalization of AS complexity into discrete categories K(*n*), where *n* refers to the theoretical number of alternative nodes and K(*n*) = 2^n^ spliced-outcomes. Schematics of K(*n*) are made up of constitutive exons (*dark gray*) and alternative exons (*light gray*) with curved lines representing all potential exon-exon junctions. **(b)**Cumulative distribution of entropy scores (i.e. entropy = Σ_*i*_Ψ_*i*_log_2_Ψ_*i*_) for simulated AS events of different categories of complexity according to (a). Cumulative distribution plots describe the proportion of data (*y-axis*) less than or equal to a specified value (*x-axis*). Cumulative Freq F(x), cumulative distribution function. See **Fig. 2b** legend for a description of cumulative distribution plots. **(c)**Comparison of the ability of different RNA-seq analysis methods to detect AS events from artificial reads (Methods) of simulated complexity as defined in (a). Bar plots show (*left*) the absolute error rate as a function of increasing complexity of AS and (*right*) the total number of AS events detected. Error bars display the standard error. Ψ, Percent Spliced In. Color legend as in panel (b). **(d)**RT-PCR analysis confirms the presence of numerous spliced isoforms in N2a cells at increasing levels of complexity matching Whippet predictions for complexity and entropy (*far right*). Boxes to right of gels display UCSC (*left, blue*) and Whippet (*right, yellow*) in silico predictions based on expected primer amplification products (Methods). Colored boxes (*blue and yellow*) represent correct predictions whereas black boxes indicate possible missed predictions. Diagrams below show exon structures of analyzed genes with approximate positions of RT-PCR primers indicated. Predicted constitutive and alternative exons are indicated in dark and light gray, respectively. **(e)**Bar plot displaying the maximum variance of splicing entropy per gene across multiple human tissues. This reveals that more than 20% of genes exhibit extensive variance in the entropy of AS across human tissues. **(f)**Functional analysis for GO, REACTOME and KEGG functional categories of genes that show large changes in splicing entropy (>2.0) across human tissues. **(g)**Symmetrical heatmap of pairwise correlations of normalized AS event entropy across multiple tissues. Heatmap showing affinity propagation clustering of pairwise similarities between splicing entropy scores. Colored bars surrounding heatmap indicate clusters defined by the dendrogram. Darker blue represents stronger correlation in splicing entropy between tissues types, whereas lighter blue indicates weak or no correlation. r1, replicate 1; r2, replicate 2 **(h)**Cumulative distribution plots comparing the cross-species variance of entropy values among the same tissue in seven vertebrates (at least three species present per-event) as compared to a permuted null control. (i) Violin plots of the distribution of splicing entropy in different cellular compartments and ribosome (mono and polysome) fractions. Kernel density is displayed as a symmetric curve, with white dots indicating the median, black box the interquartile range, and black lines the 95% confidence interval.

To assess whether Whippet can accurately quantify AS events with increasing degrees of complexity and entropy, we simulated RNA-seq datasets and corresponding Ψ values for events in these formalized categories of increasing complexity and distributed entropy (Fig. 2a-b). Importantly, in contrast to other methods, the accuracy of Whippet-derived estimates for Ψ did not decrease as the complexity and entropy of the simulated AS events increased (Fig. 2c). Moreover, in contrast to most other compared methods – with MAJIQ being the exception –, Whippet also reported a higher number of total alternative exons when analyzing increasingly complex AS events. These analyses thus indicate that Whippet offers a significant advantage over other methods in terms of its combined capacity to both detect and quantify AS events ranging from simple, independent cassette exons to complex patterns of AS involving different combinations of splice sites.

To experimentally validate Whippet-derived predictions of high AS event entropy, RNA-seq data^21^ from mouse neuroblastoma (N2a) cells were analyzed and 10 events with different predicted degrees of entropy and complexity involving tandem arrays of alternative exons were tested by RT-PCR (see Methods). Importantly, the RT-PCR assays yielded amplification products consistent with the Whippet predictions for 9/10 (90%) of the tested events (Fig. 2d, Supplemental Fig. 7). This analysis thus demonstrates a strong correlation between the AS event entropy and complexity predicted by Whippet and detected by RT-PCR assays.

### Detection of high-entropy, tissue-regulated AS events in transcripts from >20% of human genes

To investigate the prevalence and possible biological relevance of high-complexity AS events in the mammalian transcriptome, we applied Whippet to an analysis of AS across more than 60 diverse human and mouse tissue RNA-seq datasets. Remarkably, this analysis revealed that 32.97% of human (or 28.90% of mouse) protein coding genes harbor an AS event predicted by Whippet to have a high-entropy (entropy > 1.0; see Methods) event in at least one tissue. Moreover, the majority of these events are predicted to undergo large tissue-dependent changes in splicing entropy that affect the regulation of approximately 21% of genes (Fig. 2e). These regulated, high-entropy AS events display substantial expression of multiple isoforms and the corresponding genes are significantly enriched in functional annotations associated with the cytoskeleton, extracellular matrix organization, cell communication, signaling and muscle biology (Fig. 2f, p < 0.05; FDR corrected hypergeometric test). Clustering of the AS events based on pairwise comparisons of their overall entropy scores across different tissues highlighted subsets of AS events that display differences related to both tissue origin (e.g. neural- and muscle-dependent) as well as cell state (e.g. post-mitotic versus proliferative cells and tissues) (Fig. 2g, Supplemental Fig. 8a for mouse).

To further assess the possible functionality of high-entropy events detected by Whippet, we investigated their evolutionary conservation. Accordingly, Whippet was used to analyze RNA-seq data from six of the same organs from seven vertebrate species^1^ and entropy values for the orthologous exons from each species were compared. This analysis revealed a lower variance in AS event entropy values between the analyzed species than expected by chance (i.e. when compared to a randomly-permuted set of exons from the same data) (Fig. 2h; Supplemental Fig. 8 and Methods). This observation thus suggests that the entropy of AS events tends to be conserved across vertebrate species.

We next asked whether high-entropy AS events are potentially translated and thus contribute to proteomic complexity. Whippet was applied to RNA-seq data from HeLa whole cell, nuclear, and cytosolic fractions, as well as mono- and polysomes^5^. This analysis reveals comparable distributions of AS event entropy across all samples (Fig. 2i; d < 0.25, Cohen’s D statistic, Nuclear vs High Polyribosome), suggesting that high-entropy AS events contribute substantially to the translated transcriptome. Furthermore, the enrichment of high entropy events within the 5’-UTR of transcripts (Supplemental Fig. 8d, p < 4.37 × 10^−38^, Fisher’s exact test) suggests an additional role in regulating translation.

### High-entropy alternative splicing regulates genes encoding proteins with extensive domain repeats and disordered regions

Given that previous studies have illustrated the importance of AS in rewiring protein-protein interaction networks, as well as controlling additional aspects of protein function^22,23^, we hypothesized that increasing levels of AS event entropy may be associated with specific protein structural features. We observe a significant monotonic increase in the frequency of overlap with intrinsically disordered regions as a function of increasing entropy of AS events (Fig. 3a; p < 1.02 × 10^−43^, Wilcoxon Rank Sum Test, low- vs highest-entropy events). As expected, an overall inverse trend is observed for overlap with structured protein domains (Fig. 3a, p < 1.78 × 10^−41^, Wilcoxon Rank Sum Test). An interesting exception is that highest-entropy AS events (entropy > 2.0) display significant overlap with structured tandem repeat domains (Fig. 3a p < 2.14 × 10^−05^, Wilcoxon Rank Sum Test), particularly nebulin-like and epidermal growth factor (EGF)-like domains (p < 0.05, Fisher-exact test). Further analysis of the highest-entropy events (>2.0) overlapping tandem protein domain repeats reveals that they are significantly more likely to arise from exon duplication than lower-entropy events (<2.0) AS events (Fig. 3b, p < 4.57 × 10^−42^, Fisher’s exact test; Supplemental Fig. 9). As an example, Whippet detected and quantified high-entropy splicing events overlapping two classes of tandem repeat domains, LDL-receptor class A and EGF-like domains, within the low-density lipoprotein receptor-related protein 8 (LRP8). These events were confirmed by RT-PCR analysis (Supplemental Fig. 9d). Moreover, supporting the likely functional importance of AS events in this tandem array, one of them is specifically differentially regulated by the neural and muscle-enriched splicing regulator Rbfox2 (Supplemental Fig. 9d). These data support an important role for Whippet-detected, high-entropy AS events in the expansion of proteomic diversity, principally through changes to intrinsically disordered protein regions as well as through combinatorial changes to the composition of tandem arrays of specific-classes of protein domains.

**Figure 3.**
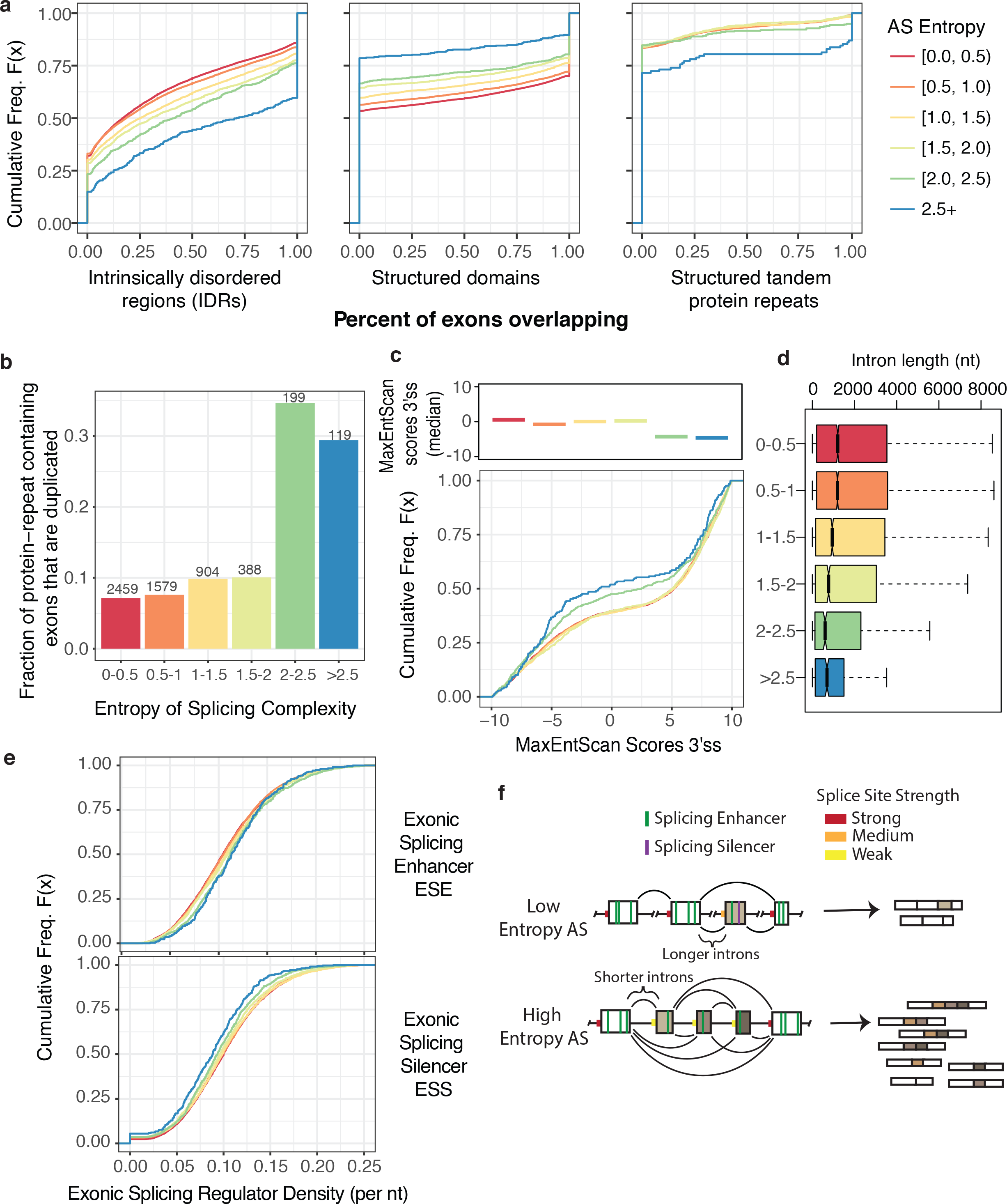
High entropy events overlap multi-domain structures and are characterized by suboptimal or reduced frequencies of cis-acting splicing elements (a) Cumulative distribution plots showing: frequency of overlap of AS events (with different degrees of entropy) within intrinsically disordered regions (IDRs) of proteins (*left*), structured single protein domains (*middle*), and structured tandemly repeated protein domains (*right*). See Fig. 2b legend for a description of cumulative distribution plots. **(b)**Bar plot showing the frequency at which exon is undergoing AS with different degrees of entropy (based on Whippet analysis of tissue RNA-seq data in Fig. 2) show evidence of duplication. Numbers of AS events analyzed indicated above plots. See **Fig. 3a** for color legend. **(c)**Plots showing the cumulative distribution of 3’-splice site (3’ss) strength estimated using MaxEntScan^48^ and binned by maximum splicing entropy scores (*bottom*). The median 3’ss strengths for AS events with different degrees of splicing entropy are plotted as colored lines (top). See Fig. 2b legend for a description of cumulative distribution plots. See **Fig 3a** for color legend. **(d)**Boxplot displaying the distribution of intron length surrounding exons binned by maximum entropy of AS. Boxplots display the interquartile range as a solid box, 1.5 times the interquartile range as vertical thin lines, the median as a horizontal line, and the confidence interval around the median as a notch. See **Fig. 3a** for color legend. nt, nucleotide. **(e)**Cumulative distribution plots showing exonic splicing regulatory elements in AS events with different degrees of AS event entropy. Scores were calculated based on density of exonic splicing enhancers (*top*) and exonic splicing silencers (*bottom*) per nucleotide. Motifs extracted from Ke et al. 2011^46^. See **Fig. 3a** for color legend. See Fig. 2b legend for a description of cumulative distribution plots. **(f)**Mechanistic model for the regulation of low-entropy (simple binary) AS events versus high-entropy (complex) AS events by *cis*-acting elements and other sequence features. Exon represented by boxes and introns by lines with *cis*-elements and relative splice site strength annotated in the legend.

### High-entropy AS events display prototypical alternative splicing signals

We hypothesized that high-entropy AS events may be associated with specific sequence features that facilitate their complex patterns of regulation. To investigate this, we binned AS events by entropy and compared the strengths of their 3’- and 5’-splice site consensus sequences, flanking intron lengths, and exonic splicing enhancer (ESE) and silencer (ESS) motif densities. Remarkably, the highest-entropy AS events show significant decreases in both 3’- and 5’-splice site strength compared to low-entropy AS events (Fig. 3c, Supplemental Fig. 10; p > 3.73 × 10^−4^ and 1.83 × 10^−3^, Wilcoxon). Additionally, we observe monotonic decreases in both flanking intron length (Fig. 3d, p < 1.78 × 10^−18^, Wilcoxon, highest vs lowest entropy events) and ESS motif density (Fig. 3e; ESS: p < 6.06 × 10^−05^; Wilcoxon, highest vs lowest entropy events) as a function of decreasing entropy. In contrast, the density of ESE elements displayed a monotonic increase between high- and low-entropy AS events (Fig. 3e; ESE: p < 4.20 × 10^−06^, Wilcoxon, highest vs lowest entropy events). These results thus suggest that weak splice sites, reduced intronic length, and altered frequencies of exonic splicing elements, are important features underlying the regulation and therefore function of high-entropy AS events (Fig. 3f).

### Whippet detects global increases in high-entropy AS in cancer samples

Aberrant pre-mRNA splicing is a hallmark of cancer and contributes to numerous aspects of tumour biology^24,25^. Cancer associated changes in AS have been linked to altered expression of numerous RNA binding proteins, some of which are oncogenic or act as tumour suppressors, as well as to splicing-sensitive disease mutations that impact the levels or activities of other cancer-associated genes^26^. Despite evidence for altered patterns of AS in cancer^27,28^, the extent to which these changes relate to altered levels of splicing complexity has not been previously determined. Accordingly, we applied Whippet to compare AS entropy using RNA-seq data (**Supplementary Table 1**) from 15 matched tumour and control liver samples of patients with hepatocellular carcinoma (HCC), the third leading cause of cancer deaths worldwide. Remarkably, this analysis revealed an overall significant and reproducible (i.e. between replicate samples) increase in AS event entropy in tumour compared to control samples (Fig. 4a-c; 3.00 × 10^−07^, Wilcoxon). Functional analysis of genes with the largest changes in entropy of AS display a striking enrichment for categories known to be dysregulated in liver cancer including DNA repair and cell-cycle regulation (Fig. 4d, p < 0.05; FDR corrected hypergeometric test). Further investigation of these events revealed a number of AS events previously identified as aberrant in cancer samples (Fig. 4e), including those associated with over-expression of the splicing regulator SRSF1^29,30^. Consistent with this observation, differential gene expression analysis revealed a number of RNA binding proteins, including SRSF1, that are significantly over-expressed in tumour compared to control samples (Fig. 4f-g and Supplemental Fig. 10c; DESeq2, False Discovery Rate-adjusted p-value < 0.001). To further investigate the possible role of SRSF1 over-expression in the expansion of AS entropy observed in cancer samples, we used Whippet to analyze RNA-seq data^29^ from an MCR-10A cell line transiently over-expressing SRSF1. These data revealed a significant increase in high-entropy AS events associated with SRSF1 over-expression (Fig. 4h; p < 9.41 × 10^−9^, Wilcoxon rank-sum test, compared to control) and a significant overlap with events differentially regulated between tumour versus normal tissues (Fig. 4i; p < 2.09×10^−5^, Fisher’s exact test). These data thus demonstrate the utility of Whippet in the detection of disease-associated changes involving high-entropy AS events and further reveal that overall splicing entropy increases in specific tumour types in response to changes in the expression of oncogenic splicing regulars, such as SRSF1.

**Figure 4.**
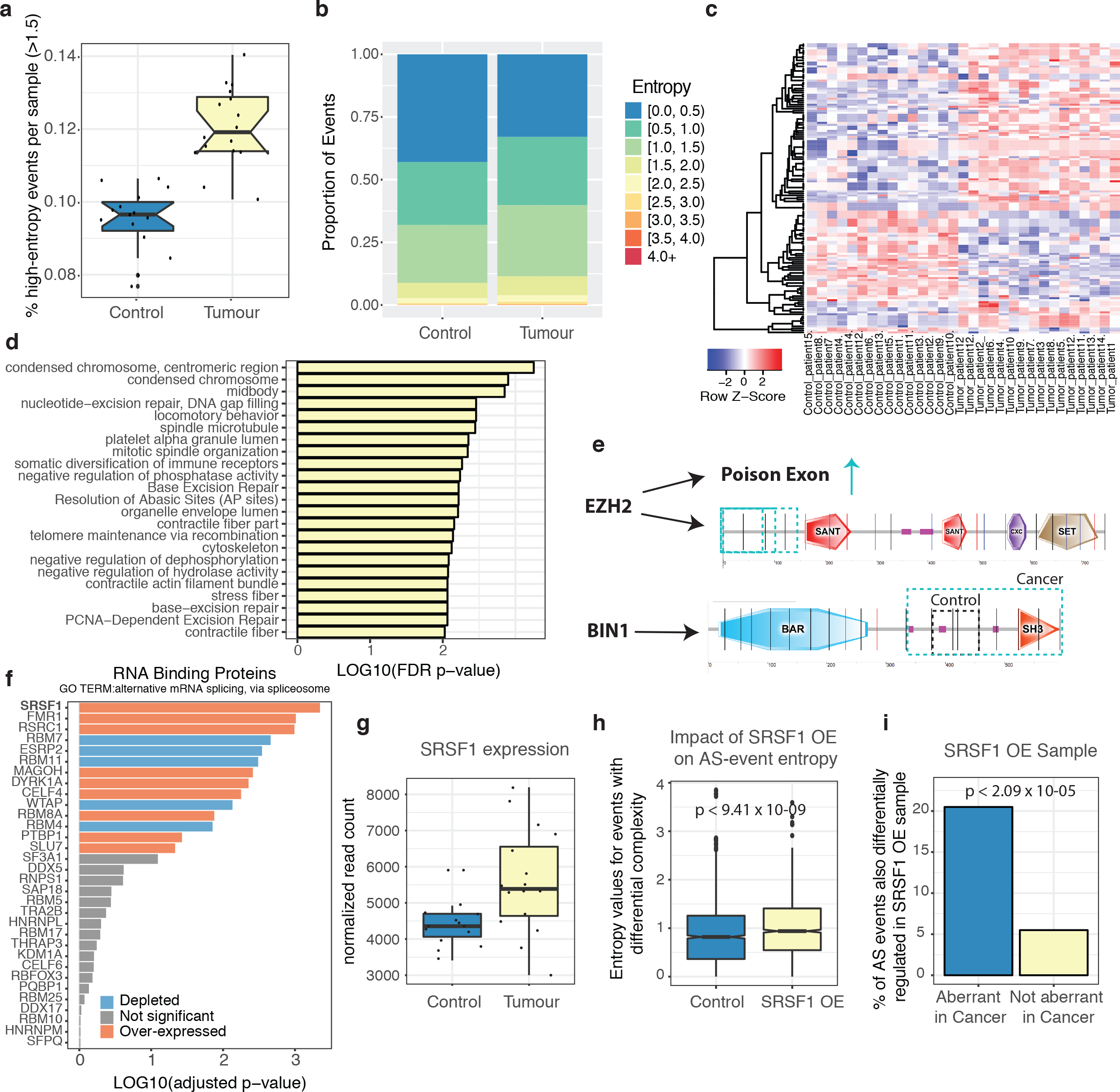
Expansion of high splicing entropy in cancer tissues associated with over-expression of the essential splicing factor, SRSF1. **(a)**Notched boxplot showing percentage of high entropy AS events (> 1.5) per sample identified from RNA-seq analysis of 15 matched tumour and control samples. Black dots represent individual data. See Fig. 3d for descriptions of boxplots. **(b)**Stacked bar plot showing the proportion of AS events of increasing splicing entropy across 15 matched tumour and control RNA-seq samples. **(c)**Clustered heatmap of splicing entropy values for differential entropy events with significant entropy changes (p>0.05, Wilcoxon rank-sum test) identified from RNAseq analysis of 15 matched tumour and control samples. **(d)**Bar plots for GO, REACTOME and KEGG functional categories of genes with events with significant entropy changes (p<0.05, Wilcoxon rank-sum test) from (c) identified from RNA-Seq analysis of 15 matched tumour and control samples. Pvalues were corrected using false discovery rate (FDR) multiple hypothesis testing correction. **(e)**Schematic diagrams of two genes showing strong and significant changes in AS event entropy between tumour sample and matched control. Domain structure extracted from SMART database ^49^. Light blue arrows and boxes indicate increased occurrence of splicing regulation in tumour samples. For BIN1, the gray box highlights protein region regulated by splicing in control samples. EZH2, Histonelysine N-methyltransferase EZH2; BIN1, Myc box-dependent-interacting protein 1 (f) DESeq2 differential gene expression analysis^50^ for selected RNA-binding proteins (GO:0000380) identified from RNA-Seq analysis of 15 matched tumour and control samples. Genes with blue bars display reduced expression in cancer samples, red bars show increased expression in cancer samples, and gray bars show no significant difference between control and tumour samples. **(g)**Boxplot showing DESeq2 normalized read counts for SRSF1. (h) Boxplot showing relative complexity of transcriptomes, as measured by distribution of entropy scores for high quality AS events, between SRSF1 overexpression (OE) sample and matched control. **(i)**Bar plot showing percentage of events from plot (h) showing differential splicing changes between SRSF1 OE (over-expression) samples and matched control that overlap with aberrant splicing changes in tumour samples from (c), as compared to expected events not identified as differential in (c).

## Discussion

In this study, we present a high-performance graph-based approach for the rapid quantification of transcriptome diversity generated by AS. Whippet applies the concept of lightweight algorithms^10,11^ to event-level splicing quantification by RNA-seq, thus enabling a significant advancement in terms of both speed and resource usage for AS analysis relative to previously published methods. Whippet also simplifies AS analysis by eliminating the requirement for extensive computational resources and user knowledge. Importantly, Whippet further affords rapid and accurate quantification of both low- and high-entropy AS events, thereby providing new insight into transcriptomic complexity.

Our results from applying Whippet indicate that high-entropy AS events occur more frequently in vertebrate transcriptomes than previously appreciated^12,31^, and further provide evidence that these events are likely biologically significant since their entropy levels are frequently tissue-regulated, conserved, and the corresponding variant transcripts are highly expressed. Many of the events are reminiscent of well-studied examples of high-entropy AS in other systems, such as the myriad of splice variants generated by tandem arrays of alternative exons in the Drosophila *DSCAM* gene^32^. In this classic example, high-entropy AS events overlap tandemly repeated immunoglobulin-like domains that function as interaction surfaces in neural circuit assembly^33^. Our results suggest that targeting of tandem protein repeat domains by high-entropy splicing represents a more widely used mechanism to modulate binding and functional specificity of multi-domain proteins. We further provide evidence that the generation of large repertoires of mRNAs from high-entropy AS events is particularly prominent in post-mitotic tissues, and likely contributes to intricate networks of regulation and cell-cell interactions in these tissues.

Alterations in RNA splicing by spliceosomal gene mutations and the over-expression of RBPs contribute to the general transcriptional dysfunction that characterizes myelodysplastic syndromes and related cancers^34^. In the present study, we demonstrate a significant increase in AS event entropy in hepatocellular carcinoma, affecting genes that function in DNA damage and spindle formation, and we relate these changes to the mis-regulation of the splicing regulator, SRSF1. These data may reflect an overall loss of splicing fidelity in cancers and exemplify how the formalization of AS entropy is important when evaluating changes in global splicing patterns^35^. For example, such measures of entropic splicing change may be valuable in future diagnostic techniques for precision medicine. Additionally, the ability of Whippet to rapidly quantify raw read data on a personal computer renders genome-wide analyses of AS accessible to a wider scientific community. In summary, the development of Whippet facilitates the efficient and comprehensive analysis of simple to complex AS events that function in normal and disease physiology.

## SUPPLEMENTAL SOFTWARE

Whippet is implemented in the high-level, high-performance dynamic programming language Julia (julialang.org) and is freely available as open-source software for academic use (Git repository: http://github.com/timbitz/Whippet.jl).

## ACKNOWLEDGEMENTS

We gratefully acknowledge M. Irimia for critical review and extensive testing of the Whippet software. We also thank M. Irimia, N. Barbosa-Morais, U. Braunschweig, S. Gueroussov, and B. Harpur, for helpful discussions and critical review of the manuscript. This work was supported by CIHR grants to B.J.B. T.S.W. was supported by CIHR and Charles H. Best Postdoctoral Fellowships. R.J.W. was supported by CIHR Postdoctoral and Marie Curie IOF fellowships. K.C.H.H. was supported by an Ontario Graduate Scholarship and a CIHR Frederick Banting and Charles Best Canada Graduate Scholarship. A.B. was supported by a CIHR Postdoctoral Fellowship. B.J.B holds the University of Toronto Banbury Chair in Medical Research.

## AUTHOR CONTRIBUTIONS

T.S.W conceived, designed, and implemented the Whippet software, with contributions from R.J.W. T.S.W., R.J.W. and K.C.H.H. simulated data and benchmarked accuracy and performance. R.J.W. and T.S.W. designed and performed computational analyses, with input from B.J.B. A.B. performed experimental validations. T.S.W, R.J.W and B.J.B. wrote the manuscript with input from the other authors.

## METHODS

### Defining and Building Contiguous Splice Graphs

The central data structure underlying *Whippet* alignment and quantification is the *Contiguous Splice Graph (CSG)*, a directed multigraph whose nodes are exonic sequence and whose edges annotate transcription start and end positions and donor and acceptor splice sites within the same gene (Supplemental Fig. 1). Briefly, a CSG separates all non-redundant exon intervals into separate ‘nodes’, and each node boundary exists as a separate edge type, allowing for various connectivity to other nodes (**Supplemental Table 2**). A *CSG Sequence* built from a given set of transcript annotations may not resemble any of the individual transcript sequences, however each of those sequences can easily be defined as a valid path or sequence of nodes leading through the graph.

Formally, a CSG contains five essential parts, an *CSG Node Vector* (*n_i_* ∈ *N*), *CSG Edge Type Vector* (*e_i_ ∈ E*, where *E* = {‘*SL*’, ‘*LS*’, ‘*LL*’, ‘*LR*’, ‘*RR*’, ‘*SR*’, ‘*RS*’}), *CSG Left Edge Vector* (*le_i_* ∈ {*I_k_ ∪ Ø*}), an *CSG Right Edge Vector* (*re_i_* ∈ {*I_k_ ∪ Ø*}) and an *CSG Sequence* (*s_i_* ∈ {*A,C,G,T*}). Here, *I_k_* represents a nucleotide k-mer of size *k* (where 0 ≤ *k* ≤ 32) encoded as an integer (*I_k_ ∈* {*0,…, 4^32^ −1=18446744073709551615*}) defined by an enumeration over the nucleotides where *nt* is a sequence of nucleotides, (*nt_i_ ∈* {*A=0,C=1,G=2,T=3*}), and *I_k_* = Σ_*i*_*nt*_*i*_,⋅2^*i*^ with *i ∈ {1,…,k}*.

Each node in *N* contains a genomic coordinate, an offset position within the *CSG Sequence* and a length in nucleotides. The edge types within set *E* are described as follows:

**Supplemetal Table 2.** Edge types and their meaning and properties. *Left* refers to the property of the *exiting* edge to the 3’ of the adjacent node to the *left*, or directly upstream of the given edge. *Right* refers to the property of the leading edge of the next node, directly adjacent and downstream of the given edge.

| Edge Type | Meaning | Splice From 5' to 3' | Splice To 5' to 3' | Extend Through | Must Splice | *Inverse*= |
| --- | --- | --- | --- | --- | --- | --- |
| ‘SL’ | TxStart (*right*) | no | no | yes | no | ‘RS’ |
| ‘LS’ | 5’ SS (*left*) & TxStart (*right*) | yes | no | no | *fwd* | ‘SR’ |
| ‘LL’ | 5’ SS (*left*) | yes | no | yes | no | ‘RR’ |
| ‘LR’ | 5’ SS (*left*) & 3’ SS (*right*) | yes | yes | yes | *fwd & rev* | ‘LR’ |
| ‘RR’ | 3’ SS (*right*) | no | yes | yes | no | ‘LL’ |
| ‘SR’ | TxEnd (*left*) & | no | yes | no | *rev* | ‘LS’ |
|  | 3' SS (*right*) |  |  |  |  |  |
| 'RS' | TxEnd (*left*) | no | no | yes | no | 'SL' |

Each edge that has the ability to *‘Splice from ’-5’to 3’*, (*e_i_ ∈ L = {‘LS’, ‘LL ’, ‘LR’}⊆E*), will have a k-mer entry (*le_i_ ∈ I_k_*) in *CSG Left Edge Vector*, while all others will be *undefined* (*le_i_ ∈Ø*). Likewise, each edge that has the ability to *‘Splice To’-5’ to 3’*(*e_i_ ∈ R = {‘LR’, ‘RR’, ‘SR’}⊆E*), will have a k-mer entry (*re_i_ ∈ I_k_*) in *SG Right Edge Vector*, while all others will be undefined (*re_i_∈Ø*).

In order to build a CSG, let *t_i_ ∈ T_g_* be the set of transcript annotations for a given gene *g*, and let *x_j_ ∈ X_g_* where *X_g_ = ∪_i=1,…,g_ X_i_* be the full set of exon (start, end) intervals *x_j_* = (*a_j_, b_j_*) from all transcript annotations in gene *g* where *a_j_ ≤ b_j_* and *a_j_ ∈ A_g_* and *b_j_ ∈ B_g_*. We define *X_g_* as an ordered and enumerated set where a given exon interval is less than or equal to the next exon interval *x_j_ ≤x_j+1_* satisfying the boolean (*a_j_* < *a_j+1_*) ∨ ((*a_j_* == *a_j+1_*) ∧ (*b_j_* ≤ *b_j+1_*)) regardless of transcript strand. Further, we define a single ordered vector of exon start or end positions *ν_i_* ∈(*V_i_*={*A_g_ ∪ B_g_*}) where *ν_i_ ≤ ν_i+1_*. A contiguous splice graph is built, *csg_g_ ∈ G*, by iterating through *ν_i_*. The pseudocode is given below (**Algo. 1**):

#### ‘‘‘Algorithm 1.

~~~
Function *stranded_push!*( list, element ):
       If *strand* of *t_i_* is *positive*:
                *push!* ( list, elem )
       Else:
                *unshift!* ( list, elem )

Function *stranded_pop!* ( list, element ):
       If *strand* of *t_i_* is *positive:*
                *pop!* (list, elem)
       Else:
                *shift!* (list, elem)
*e* = [‘SL’]
For *ν_i_* in *{A_g_ ∪ B_g_}:*
 If *X_g_ contains subinterval (ν_i_, ν_i+1_*):
       *stranded_push!*(*n, SGNode(ν_i_, ν_i+1_*))
       *stranded_push!*(*s, SGSequence(ν_i_, ν_i+1_*))
       (*ν_i+1_ ∈ A_g_*) ∧ *stranded_push!(e, ‘LL’*)
       (*ν_i+1_ ∈ B_g_*) ∧ *stranded_push!(e, ‘RR’*)
       (*ν_i+1_∈ F_g_*) ∧ *stranded_push!(e, ‘SL’)*
       (*ν_i+1_ ∈ L_g_*) ∧ *stranded_push!(e, ‘RS’*)
 Else:
       *stranded_pop!*
       (*ν_i_ ∈ B_g_ ∧ ν_i+1_ ∈ A_g_*) ∧ *stranded_push!(e*, ‘LR’)
       (ν_i_ ∈ L_g_ ∧ ν_i+1_ ∈ A_g_) ∧ stranded_push!(e, ‘SR’)
       (*ν_i_ ∈ B_g_ ∧ ν_i+1_ ∈ F_g_*) ∧ *stranded_push!(e, ‘LS’*)
 If *strand* of *ti* is *positive*:
     *s = reverse_complement!(s)*
     *e* = *inverse*(*e*)
’’’
~~~

### Contiguous Splice Graph Alignment

After building all CSGs, the *CSG Sequences* are concatenated into a single *Multi-CSG Sequence* that is used to create a non-redundant transcriptome FM-Index^13^ for compressed full-text substring search. Since a major computational bottleneck of formal read alignment is the full-text search for alignment seed location, we focused on making the entire seeding process as efficient as possible. Whippet therefore will not attempt to seed any window containing a FASTQ nucleotide entry with a QUAL score below the minimum threshold (Phred Q=4, P=0.398 by default), and will iteratively increment until a seed of acceptable QUAL distribution is obtained. After choosing a seed of (constant seed-length *n*) at read offset *i* (S_*i,n*_), Whippet is very restrictive in the number of matching loci (Count(S_*i,n*_)) that a valid seed is allowed to have (by default 1 <= Count(S_*i,n*_) <= 4) in order to avoid exhaustive searching in highly repetitive sequence space. For failed seeds, the reverse complement of the seed is tried S_*i,n*_= RevComp(S_*i,n*_), and then if that fails, the seed is incremented a constant number of nucleotides (**Algo. 2**). For paired-end sequencing reads, the same principles apply, except that mate-pair seeds (M_*j,n*_) are locked into a relative orientation to one another (*fwd_read & rev_mate* by default) and only loci within the mate-pair mapping distance where || Location(S_*i,n*_) – Location(M_*j,n*_) || <= *MaxMateDistance*, are returned.

#### ‘‘‘Algorithm 2.

~~~
*seed_tries* = 0
While *i+n-1* < = length(Read) && *seed_tries < = MaxSeeds*:
                If minimum(*Qual_i,…,j+n-1_* ) > *MinQual:*
                      *seed_tries = seed_tries + 1*
                      If 1 <= Count(S_*i,n*_)<= 4:
                            *return* Locations(S_*i,n*_), *positive_strand*
                      Elseif 1 <= Count( RevComp(S_i,n_)) <= 4:
                            *return* Locations(RevComp(S_*i,n*_)), *negative_strand*
                *i* = *i* + *SeedIncrement*
~~~

Given a set of mapping transcriptome loci for a sequencing read, Whippet performs ungapped extension of each alignment seed, storing only the offset of the alignment in the read (*r*) and transcriptome (*k*), and the alignment path (*A*) through the CSG. An alignment path is defined as the vector of nodes, such that each node records the {Gene, Node, Score} = {*g, n, s*}, where the Score refers to a set of {Matches, Mismatches, and the Mismatch_probability_sum} = {m, x, p}. Alignments are extended first in the *forward* and then in the *reverse* direction from the seed offset. Alignment extensions continue until the edge of the read, a non-extendable edge (‘SR’ for *forward* direction, and ‘LS’ in *reverse* direction), the end of the CSG, or the mismatch_sum *p* exceeds the *MismatchThreshold* (default 2.0). An alignment is considered valid if its score exceeds the *MinimumScoreThreshold* (default = 75% of read contains *matches*) or contains at least one spliced edge in its path. Upon alignment extension into an unspliced node boundary (not a ‘Must Splice’) that has an ‘Extend Through’ property (**Supplemental Table 2**), the extension traverses past the boundary without splicing into the neighbor node (Supplemental Fig. 2). If the alignment ends in the neighbor node within a specified distance (k-mer size) from the node’s boundary, then the neighbor node is removed from the path, and k-mer spliced extension is attempted. If multiple unspliced boundaries are traversed before the alignment fails, spliced extension is attempted on each previous neighbor node. For spliced extension, the sorted list of (gene, node) pairs for the k-mer upstream of the node boundary, is intersected with the sorted list of (gene, node) pairs for the next adjacent k-mer (if sufficient read length exists). Compatible (gene, node) pairs for intersection must share the same gene, where the 3’ splice-site node lies downstream of the current node in the CSG (unless circular splicing flag is enabled). Alignment extension continues to all compatible downstream nodes recursively, returning only the best scoring alignment path.

### TPM Quantification and Multi-mapping Reads

For each alignment seed, an alignment is returned. Multi-mapping reads with multiple *valid* alignments whose scores are within the *AlignmentScoreThreshold* distance from the maximal scoring alignment are subsequently treated as *repetitive alignments.* Since multi-mapping reads suggest that a sequencing read could have derived from one of multiple paralogous gene loci, we utilize the Expectation Maximization (EM) algorithm to iteratively maximize the likelihood of the relative abundance of all CSGs in set G. The EM algorithm alternates between estimating the fraction of multi-mapping read counts belonging to each CSG (E-step) and calculating the relative abundance of all CSGs given the total and partial counts allocated to each gene (M-step) ^36^. In the first step, the probability of observing reads from a given CSG *i* with relative expression level *μ*_*i*_ is given by 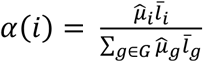
where 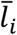 refers to the mean (effective) length of the *N* annotated transcripts *t ∈ T_i_* used to build CSG _*i*_ given a constant read length *m*, where 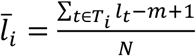. We then define a multi-mapping compatibility matrix *y_r,i_* = 1 for a read *r* that maps to a CSG *i* and *y_r,i_* = 0 otherwise ^11^. The probability of observing a specific multi-mapped read *r* from a CSG *i* in its compatible set of CSGs is then estimated in the (E-step) of the algorithm by:

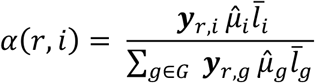

In the (M-step) of the algorithm, the relative abundance of each CSG is calculated by summing all of the full and partial read counts for each CSG *i*:

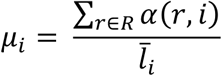

The transcripts-per-million^37^ of each CSG is then calculated as:

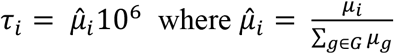

To seed the EM algorithm, a uniform probability across compatible CSGs is assigned for each read, followed by the standard M-step, and subsequent EM-steps until the end condition, (τ_*i,iter*_ – τ*_i,iter-i_*) < 0.1, or a user-defined max-iterations is reached.

### Local Splicing Event (LSE) Definition and PSI Quantification

After all reads have been assigned full or partial counts to the CSGs, it is necessary to build local splicing event (LSE) structures to quantify AS. In order to define an LSE, the set of edges connecting to, and spanning over the target node (*n*) are collected (where the read count of a spanning edge must be >= 1% of the maximal connecting edge read count). Subsequently for each node within the upstream and downstream boundary nodes, we also collect the connecting and spanning edges, extending the upstream and downstream boundary nodes as necessary to encompass all relevant edges. After collecting all edges for an event *E* as a vector of connected nodes, where *E_i_* refers to one edge in the LSE, we build a vector *V* that contains the minimum set of non-redundant paths (each path *V_i_* contains the set of nodes in the connected path) through the LSE using the edges in *E*. To build the set of paths *V* through the LSE, a recursive stitching algorithm is defined (see **Algo. 3**).

#### ’’’Algorithm 3

Function has_terminal_overlap( a, b ):

~~~
        *return* first(a) == last(b) OR first(b) == last(a) ? *True* : *False*
~~~

Function build_paths(*E, V*=copy!(*E*)):

~~~
         *R* = Vector(Ø)
         *i* = 1
         While *R* does not equal *V*:
               If *i* > 1:
                       *V* = *R*
                       *R* = Vector( Ø)
               For *j* in 1 to length(*V*):
                        Added = False
                        For *k* in 1 to length(*E*):
                                  If has_terminal_overlap( *V_j_, E_k_* ):
                                          Added = True
                                          push!( R, *V_j_ ∪ E_k_* )

                        Unless Added is True:
                               push!(R, *V_j_*)
                     *i* = *i* + 1
     *return R*
’’’
~~~

In order to quantify the paths *i ∈I* in an LSE, we utilize the set of edges *e ∈E* in the LSE and the counts *α* for some edge *e*, *α*(*e*). Counts for each unique edge *e* that exist in only one path *i* are assigned fully, *α*(*e*,*i*) = *α*(*e*). Non-unique edges existing in multiple paths have counts initially divided among their compatible paths with uniform probability, and then the maximum likelihood for the relative expression of each LSE path is estimated using the EM algorithm. We define a compatibility matrix y_*e,i*_ = 1 for an edge *e* existing in a path *i*, and y_*e,i*_ = 0 otherwise. In the M-step, the relative expression of each LSE path (*ψ_i_*) is given by:

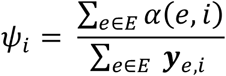

In the E-step, the counts *α*(*e,i*) for each edge *e* are divided among path *i* following:

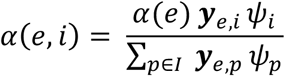

The percent-spliced-in Ψ of the node *n* is then calculated as the sum of the normalized relative expression of the paths containing the node *{I_n_ ⊂I}*:

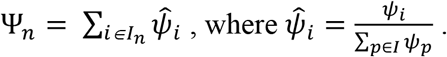

Since the EM-algorithm provides only a point-estimate for Ψ without a depth-dependent measure of variance, we utilize the conjugate posterior distribution of the binomial likelihood as a means to compute a confidence interval over Ψ. Given a total read depth for an LSE of *N* reads which can either support inclusion of node *n*, *inc ∈I_n_*, or support exclusion, *exc ∈{I – I_n_}*, the number of inclusion reads *N_inc_* are binomially distributed such that *N_inc_∼Binomial(n=N*, p=Ψ). Given a uniform prior distribution of P(Ψ) = *Beta*(*α*=l, β=l), we obtain a posterior distribution, P(Ψ|*N_inc_*) ∝ P(*N_inc_*|Ψ)P(Ψ), where P(Ψ|*N_inc_*) = *Beta*(*N_inc_* + *α*,*N_exc_* + β). A 90% confidence interval (between 5% and 95%) is then calculated through the quantile distribution of the posterior.

### Event Types and Standard Output

In order to define the nature of an alternative node in an LSE, a number of discrete categories are utilized. These include AS specific types for alternative 5’ or 3’ splice-sites, Core-exon nodes (which may be a whole exon or part of an exon with flanking alternative splice sites that are used), or a retained intron. Additionally, alternative 5’ or 3’ ends are annotated as either alternative first or last exons, or tandem 5’ or 3’ untranslated regions (UTR). **Supplemental Table 3** provides a list of the two-letter symbols for each node type and their formal definition as a set of flanking edge types.

**Supplemental Table 3.**
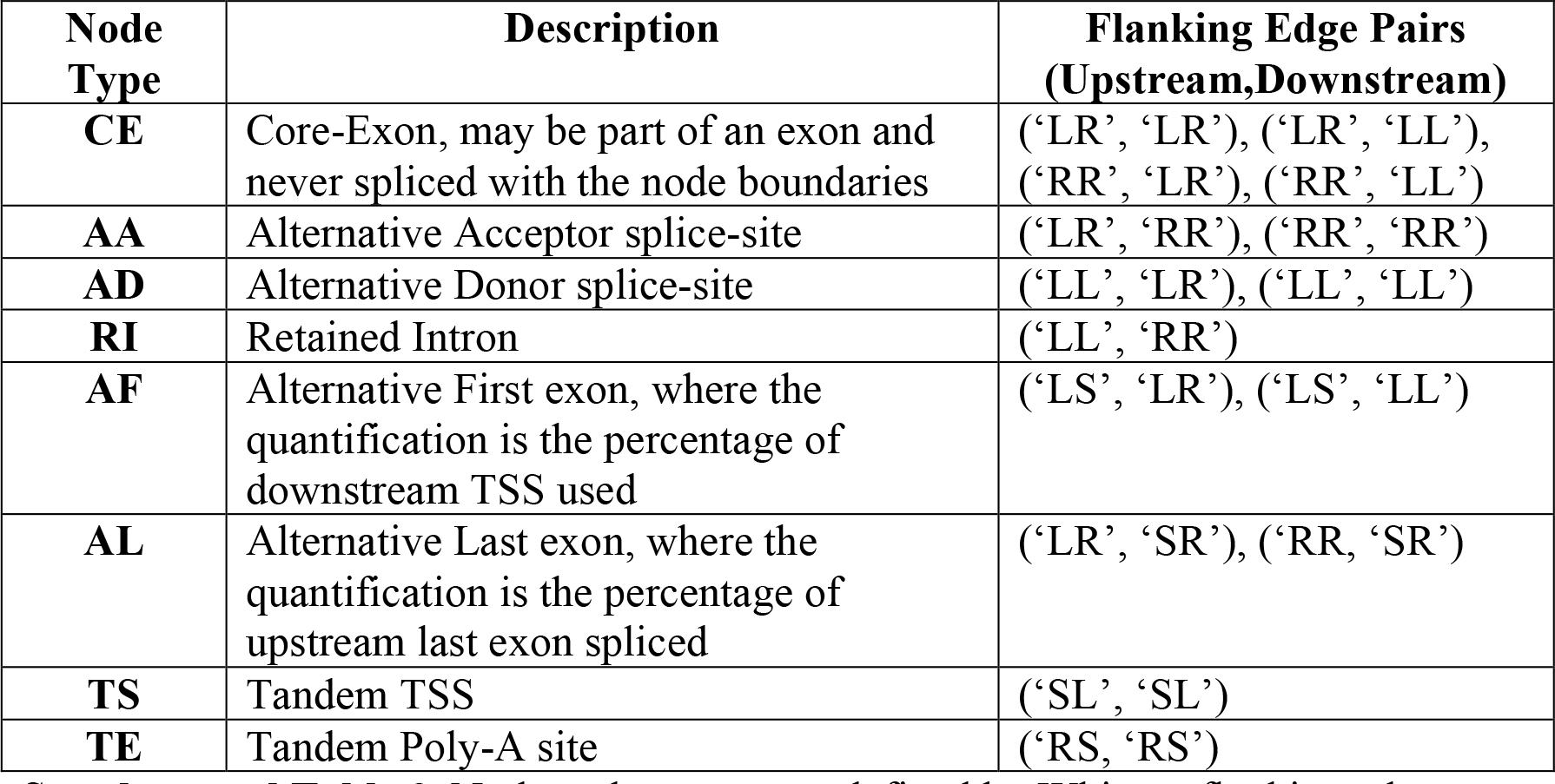
Node and event types defined by Whippet flanking edges.

### LSE Complexity and RNA-seq Simulation

In order to investigate the accuracy of AS quantification tools at higher-levels of AS complexity, it was necessary to simulate transcripts with AS-events of increasing complexity. Formalizing AS-events into discrete classes of complexity K(*n*) = 2^n^ splicing-outcomes for K1 through K6, we randomly choose 500 CSGs for each complexity class with at least *n* total internal nodes (not including nodes with transcription start or end annotations). From those CSGs, we randomly choose a set of *n* consecutive internal nodes and created partial transcript sequences from the first internal node until the last internal node with all combinations of those *n* internal nodes. In the case of alternative 5’ or 3’ splice-site nodes (AA and AD types) or retained-intron (RI type) nodes, less than 2^*n*^ total combinations were created, as the AA and AD nodes cannot be included in the transcript without all AA nodes downstream of an AA-node until its core-exon CE node, or all the AD-nodes upstream and its associated CE-node. Similarly, a retained-intron node can only be included in a transcript if both the upstream and downstream CE nodes are included.

Given the eight sets of simulated events of complexity K(*n*) (where *n* = 1,…, 6), we used polyester^38^ to simulate RNA-seq reads from the simulated transcripts for each gene. To simulate AS-events with known Ψ-values, we randomly subsampled one of the *n* alternative nodes from each gene and assigned it a random Ψ-value sampled from a Beta distribution. At higher complexities, the assignment of a Ψ-value to one node has indirect effects on the Ψ-values of the neighboring *n* alternative nodes. In order to achieve a near uniform coverage of total Ψ-values for each complexity K(*n*), the *α* and β parameters of the Beta distribution were selected *ad hoc* accordingly. For K <= 2, we used Beta(*α*=0.9, β=0.9) which produces a near uniform distribution over Ψ. For K(*n*) in the range of interval *n* ∈ [3, 5], it was necessary to use a distribution skewed more towards 0.0 and 1.0 (such as Beta(*α*=0.7, β=0.7)), since a uniform distribution of initial Ψ-values resulted in a bell-shaped curve centered on Ψ=0.5. For K >= 6, this effect was substantially increased, requiring a more skewed initial distribution, Beta(*α*=0.2, β=0.2). Transcripts containing the sampled exon were randomly assigned relative expression values such that their total expression would be proportional to the pre-assigned Ψ-value. Similarly, the remaining transcripts were randomly assigned expression values such that their total expression is proportional to (1 – Ψ). To simulate differential gene expression, each gene was randomly assigned a coverage multiplier value from a uniform distribution between 5 to 60x. Subsequently RNA-seq reads of length 100nt were simulated for each transcript in both single and paired-end modes.

### Sequence annotation

All genomic and transcriptomic sequences, as well as gene transfer format (GTF) files, were downloaded from the Ensembl database^39^. Exon annotations (including genomic annotations) were downloaded from Ensembl using BioMart^39^. The following genome builds were used: Hg19 and Mm10 using only junctions within transcripts that have a transcript support level (TSL) of 2 or less.

### Benchmarking

All benchmarking has performed on a Sun Microsystem X4600M2 server with 8 AMD Dual-Core 8218 CPU @2.6GHz, total 16 cores and 64GB RAM. Local hard disk was SATA 73GB, 10K RPM. Identical paired-end HeLa data of increasing read-depths were used for all resource usage benchmarking (see **Supplementary Table 1**). All programs were run with default settings with additional settings described in Supplemental Table 4. Resource usage benchmarks for STAR and TOPHAT included conversion to a sorted BAM file (as this step is required for majority of splicing quantification algorithms) whereas quantification time was removed for Whippet in comparison to alignment programs. When necessary, initial read alignment was done by STAR and same BAM output file used by all splicing quantification algorithms. The default linux package “time” was used to measure the resource usage of each program. The running time was calculated by combining the User time and the System time.

Benchmarks of the mapping success used wgsim (https://github.com/lh3/wgsim) simulated reads based on Hg19 GRCh37.73 Ensembl transcriptome data with error rates of 0.005 and 0.01, respectively and a read depth of ∼50M. Identical parameters were used for resource and mapping benchmarks with default parameters used (see below for details). For Whippet mapping success only considered reads mapping over splice junctions by at least 9nt.

RT-PCR values and matched RNA-seq datasets were extracted from datasets provided by Vaquero-Garcia et al. 2016 ^12^. ΔPSI calculated by comparing PSI values across cerebellum and liver samples.

**Supplemental Table 4.**
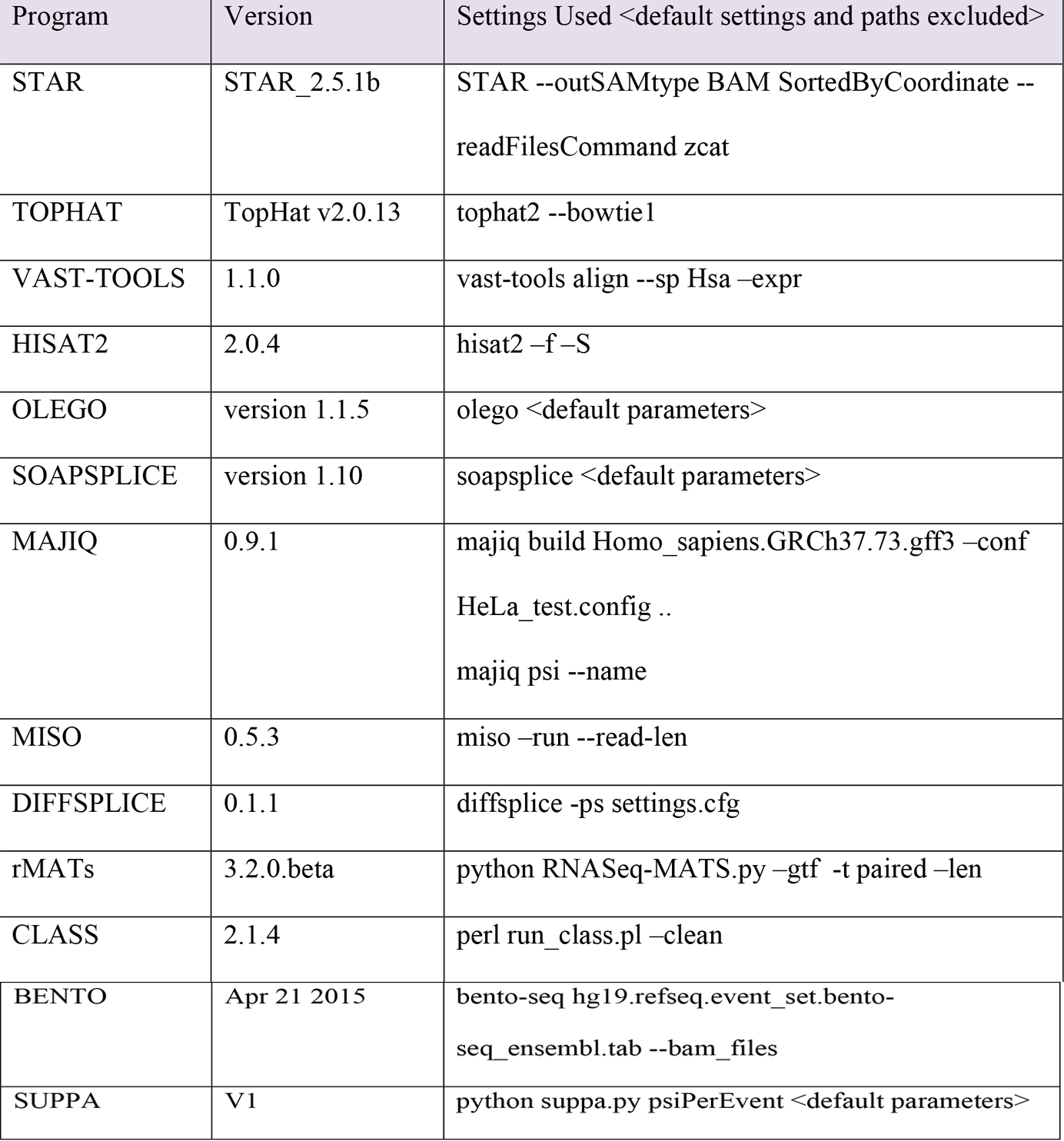
Versions of programs used in benchmarking with parameters used for alignment and splicing quantification. Default settings, paths, Fastq files excluced.

### High entropy categories and differential entropy

High entropy events are defined as entropy events with a score of greater than 1.0, differential entropy requires a change of entropy of greater than 1.0 (unless stated). Highest entropy events are greater than 2.0.

Only events with at least a read count over 30, and Ψ scores of over 0.05 and under 0.95 were included in the analyses. Analyses limited to cassette, first and last exon events (as defined by Whippet).

### Tissue-wide analysis of splicing

Tissue data was extracted from Illumina Bodymap2 dataset and supplemented by human tissue samples from Kunming Institute of Zoology (See **Supplementary Table 1**). The maximum change in splicing entropy between tissues is the comparison of the lowest entropy of an exon/node compared to the highest entropy for same exon/node across tissues. This is therefore not a measure of tissue-specificity but rather a measure of maximum variability for the number of well-expressed exon-exon junctions an exon may have across tissues.

### Polysome analysis of entropy

Monosome and polysome samples combined based on sub-groups identified within original paper ^5^. This included 80S (monosomes), low polysomes (two-four ribosomes), high polysomes (five-eight+ ribosomes), and total cytoplasmic RNA. Additional nuclear and whole-cell HeLa fractions originating from a different paper were also analysed as a comparison.

### Functional analysis

Functional analysis was undertaken using the functional enrichment analysis tool g:Profiler ^40^. Genes identified as containing mammalian-classifying splicing events were compared to a background of multi-exon genes conserved within vertebrates. Structured controlled vocabularies from Gene Ontology organization, as well as information from the curated KEGG and Reactome databases were included in the analysis. Only functional categorizes with more than 5 members and fewer than 2,000 members were included in the analysis. Significance was assessed using the hypergeometric test with the multiple testing correction method created by Benjamini and Hochberg.

### Protein feature analysis

For all positions in a protein a score for intrinsic disorder is computed using IUPred^41^. Amino acid residues with a score larger than 0.4 are considered disordered. For each coding exon the ratio of disordered residues is estimated.

For all positions in a protein low complexity regions were calculated using SEG^42^. Only amino acids not within ordered Pfam annotated protein domains^43^, putative transmembrane domains, signal peptides and coiled coil regions were considered as low complexity regions. For each exon, the ratio of amino acids annotated as within a low complexity region is estimated. Tandem protein repeat regions within structured regions were identified using the PTRStalker algorithm for de-novo detection of fuzzy tandem repeats^44^ and filtered using IUPred (score < 0.4).

### Exon Duplication

Exon duplication events were identified using approach described by Letunic et al. (2002)^45^. In brief, exon were considered duplicates if (i) within the same gene body (ii) blastn comparison had an e-value of less than 0.0001 (iii) 80% similarity in length.

Exons with peptide repeats were extracted and maximum entropy value (across tissues) identified. For each entropy bin, the fraction of duplicated exons was calculated.

### Splicing Motifs

MaxEntScan was used to estimate the splice site strength of both the 3’splice sites and 5’ splice sites. As recommended by developers, the 5’splice site strength was assessed using a sequence consisting of 3nt within exon and 6nt of intron. Similarly, the 3’splice site strength was assessed using a sequence consisting of 20nt of intron and 3nt of exon. Exonic splicing silencer or exonic splicing enhancer densities were extracted from designated motifs as quantified by Ke et al.^46^. To calculate exonic splicing enhancer/silencer density, all motifs defined by Ke et al. were summed together and normalized by number of nucleotides in exon.

### Cancer Data and SRSF1

For Figures 4A and B, all events passing the aforementioned criteria (see high entropy categories and differential entropy) were included in analysis. Differential complexity between control and tumour samples across 15 replicates (dependent on read coverage) described in Figure 4C was determined by a Wilcoxon rank-sum test (p < 0.05) of entropy scores. Differential gene expression was calculated using DESeq2 (adjusted p-value < 0.05). SRSF1 over-expression data extracted from Anczukow et al. and processed by Whippet only events with high entropy (> 1.5) in either control or over-expression study included in analysis. Events considered aberrant in splicing in Fig. 4i are displayed in Fig. 4c.

### Clustering

Unless stated all heatmaps were constructed using affinity propagation clustering with pairwise similarities as correlations and negative correlations taken into account ^47^.

### RT-PCR

Predictions on size of bands were made using UCSC server and by combining exons together from Whippet predictions. Only predictions supported by multiple sources included in figures (i.e. PCR, Whippet or UCSC).

### Slmap

Forward: GAGCGCACTCAGGAAGAGTT ; Reverse:

TTCCTTTGCTTTTGCCTGAT

### Eps15l1

Forward: TTGGAACCCTAGACCCCTTT ; Reverse:

CTTTTTCACTCTCCCGCTTG

### Asap1

Forward: GCCCGCGATGGAATAATG ; Reverse:

TGAGGAAGAGGCACAGGTCT

### Eml4

Forward: TCCTGTATAACCAATGGAAGTGG ; Reverse:

CATTGTAATTGGCCGACCTC

### Atp8a1

Forward: CGGTCGTTACACAACACTGG ; Reverse:

GGCCAAGTTCCTCATTCAGA

### Sfl1

Forward: TCATGCCTCACAAAACTGGA ; Reverse:

CCATAGCCAGCCTCTGTACC

### Mapt

Forward: AATGGAAGACCATGCTGGAG ; Reverse:

GCCACACTTGGAGGTCACTT

### Lrp8

Forward: CGGAGAGAAGGACTGTGAGG ; Reverse:

CAGTGCAGATGTGGGAACAG

### Gtf2ird1

Forward: CCCCAACACCTATGACATCC ; Reverse:

CGCTTGGGAATGTTGTCTTT

### Rbms3

Forward: GAGACAGGGTCAGAGCAAGC ; Reverse:

AAACCGGAGGCCAACTAACT

### Cask

Forward: AGGGAAATGCGAGGGAGTAT ; Reverse:

GTCATCCTTGGCTGGATCAT

### Slmap (Control)

Forward: GAGCGCACTCAGGAAGAGTT ; Reverse: TTCCTGCTCAGTCATTTCAAAC

### siRNA knockdown & RT-PCR

Mouse Neuro2A (N2A) cells were transfected with SMARTpool siRNAs (Dharmacon) (50nM final concentration), using Lipofectamine RNAiMAX (Invitrogen) as recommended by the manufacturer. A non-targeting siRNA pool (siNT) was used as a control. Cells were harvested at 48 hours post transfection and total RNA was extracted using RNeasy columns (QIAGEN). Semi-quantitative RT-PCR was performed using the QIAGEN One-Step RT-PCR kit as per the manufacturer’s instructions, using 50ng total RNA as template in a 20uL reaction and resolved on 2-4% agarose gels. Bands were quantified using Image Lab (BioRad) or ImageJ.

### Statistical Tests

In functional analysis the significance was assessed using the hypergeometric test with the multiple testing correction method created by Benjamini and Hochberg. Wilcoxon rank-sum test non-parametric statistical tests were used for comparing distributions. Affinity propagation clustering with either pairwise similarities as correlations (Pearson) or mutual pairwise similarities of data vectors as negative Euclidean distance was used to create clustered heatmaps.

## Supplementary Legends

**Supplementary Figure 1.**
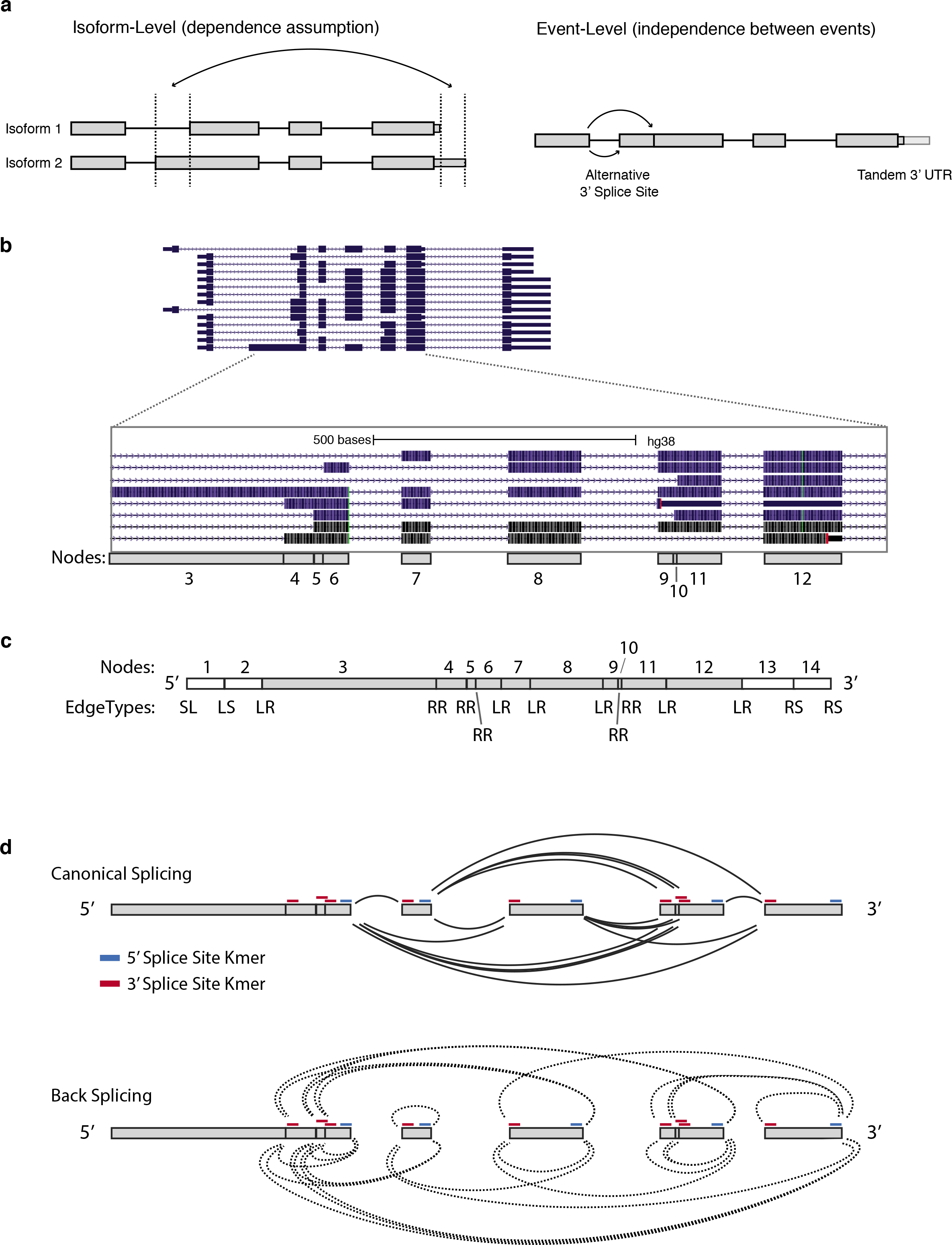
**(a)** Schematic of isoform-level (*left*) vs. event-level (*right*) quantification paradigms. An isoform-level approach to quantification with limited number of annotations, assumes dependence between transcript features, whereas the event-level approach assumes independence and measures the expression of each feature separately. **(b)** Exemplar gene and isoform annotations with a zoomed-in view of a region with complex AS patterns. The node diagram below displays how such a complex set of splicing patterns can be collapsed into a set of nodes for a CSG. **(c)** Diagram of the edge types associated with the node set in panel (b) (see **Supplemental Table 2** for description of edge types). **(d)** An overview of the possible spliced edges in the CSG, both forwards (*solid lines*) for linear products and backwards (*dotted lines*) for potentially circular products (provided as an optional command line parameter ‘–circ’).

**Supplementary Figure 2.**
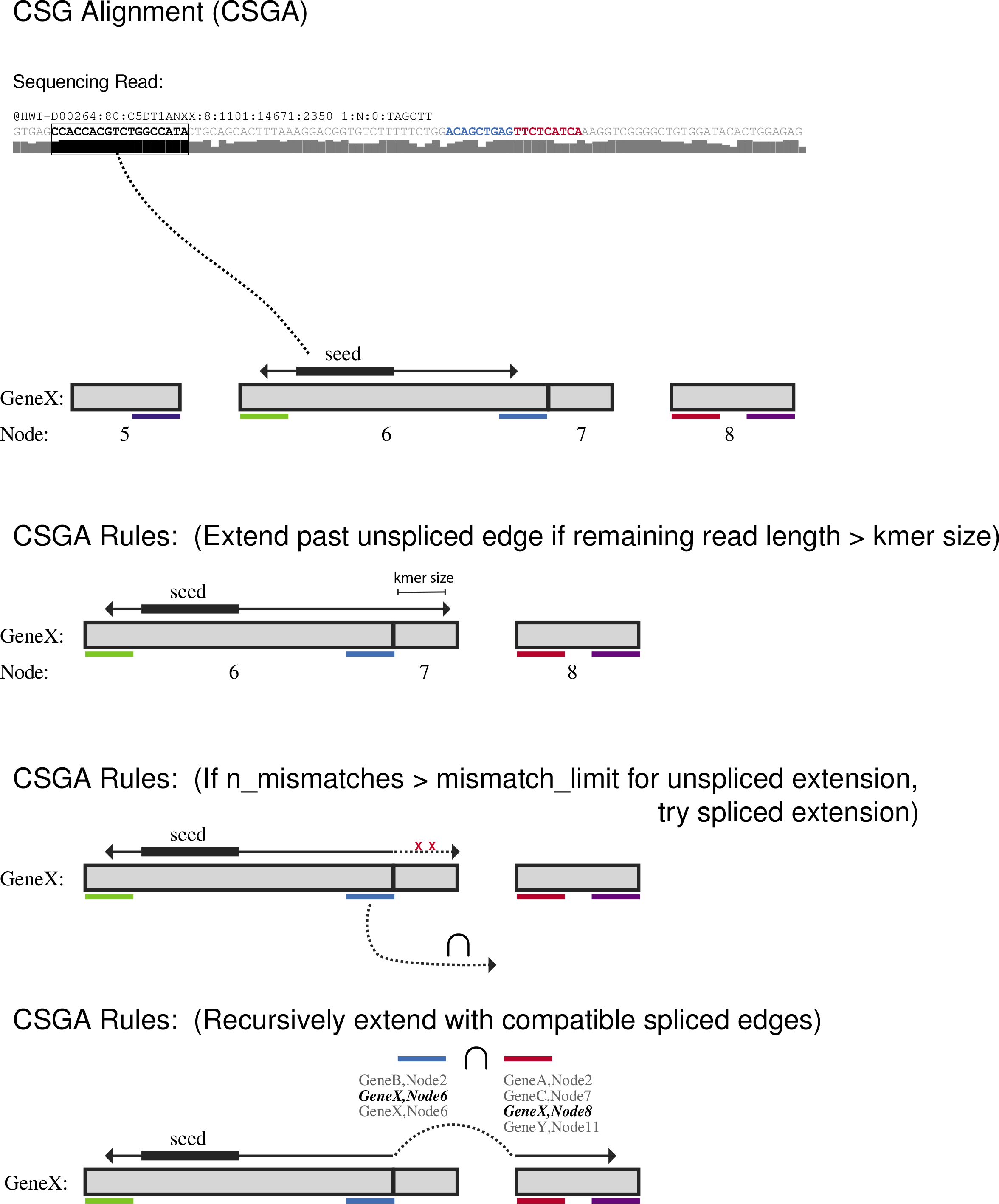
Graphical overview of the CSG Alignment algorithm. High FASTQ-QUAL region (*black box*) of sequencing reads are used to seed to the transcriptome. Alignment extension occurs in the forward and reverse directions, and spliced extension is allowed as necessary to bridge spliced edges. Unspliced alignment extension can occur past an edge that allows both unspliced and spliced extension. If the alignment fails, then the new node is removed and spliced extension proceeds from the previous node(s) (see Methods).

**Supplementary Figure 3.**
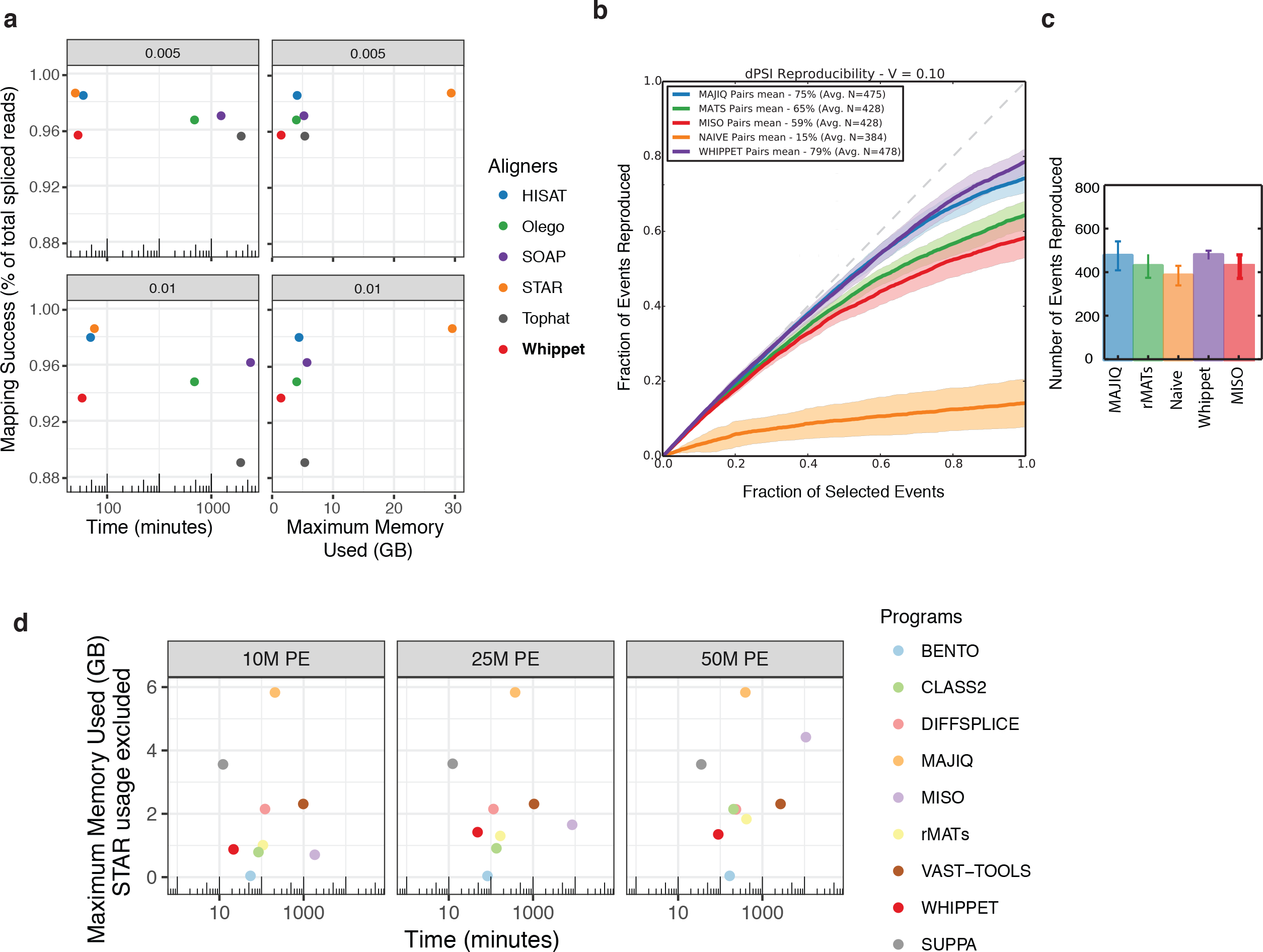
**(a)** Comparison of the mapping success (*y-axis*) and (left) log-scaled time (*x-axis*) and (right) maximum memory usage (*x-axis*) requirements of Whippet relative to several published methods for RNA-seq read alignment. Reads were simulated using wgsim (https://github.com/lh3/wgsim) with base error rate of 0.005 (*top*) and 0.01 (*bottom*). Only simulated reads overlapping exon-exon junctions by at least Whippet’s k-mer size are considered in mapping success. **(b)** Plot highlights the reproducibility of PSI values when comparing RNA-seq from two conditions. A differentially included event is considered replicated if it maintains a rank at least as high as *N* in biological replicates, where N is the set size. (see Vaquero-Garcia et al. 2016^12^ for details). **(c)** A bar chart shows the number of AS events identified by each method and used in (b). **(d)** Extension of the bottom panel in Figure 1e to include comparison with the splicing algorithm *SUPPA*. The panels compare resources used by Whippet and eight other splicing quantification methods when analyzing aligned reads (10M, 25M or 50M) (see Methods). With the exception of Whippet, SUPPA and VAST-TOOLS, all algorithms require initial alignment of reads by an aligner, in this case STAR. Resource usage of SUPPA includes initial alignment by the transcript-level expression quantification algorithm Kallisto^11^. GB, gigabyte.

**Supplementary Figure 4.**
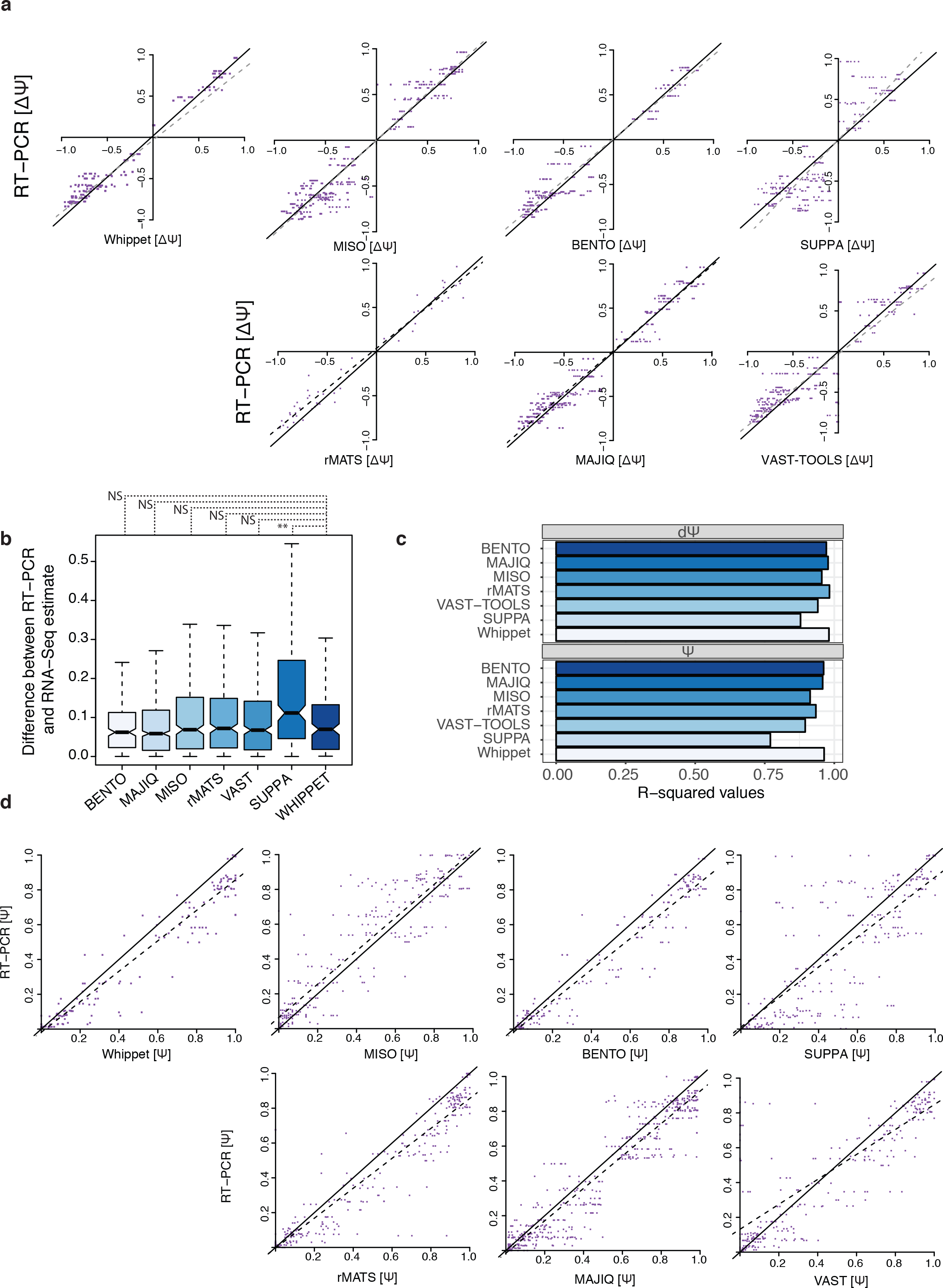
**(a)** Extension of Fig. 1d. Comparison of Whippet and six other state-of-the art published splicing algorithms for Ψ predictions from RNA-seq data to coupled Ψ assessments from RT-PCR. ΔΨ (change in percent-spliced-in) measures correspondence between liver and cerebellum samples. Regression line is shown as dotted line whereas diagonal is solid line. **(b)** Boxplots quantifying differences in Ψ between predictions from RNA-seq data and coupled Ψ assessments from RT-PCR. **(c)** R-squared values of correlations between predictions from RNAseq data and coupled Ψ assessments from RT-PCR. **(d)**Same data as in (a) but showing absolute Ψ rather than ΔΨ. Ψ, percent spliced in; NS, not significant; **, p < 1.0 × 10^−05^, Wilcoxon rank sum one sided test.

**Supplementary Figure 5.**
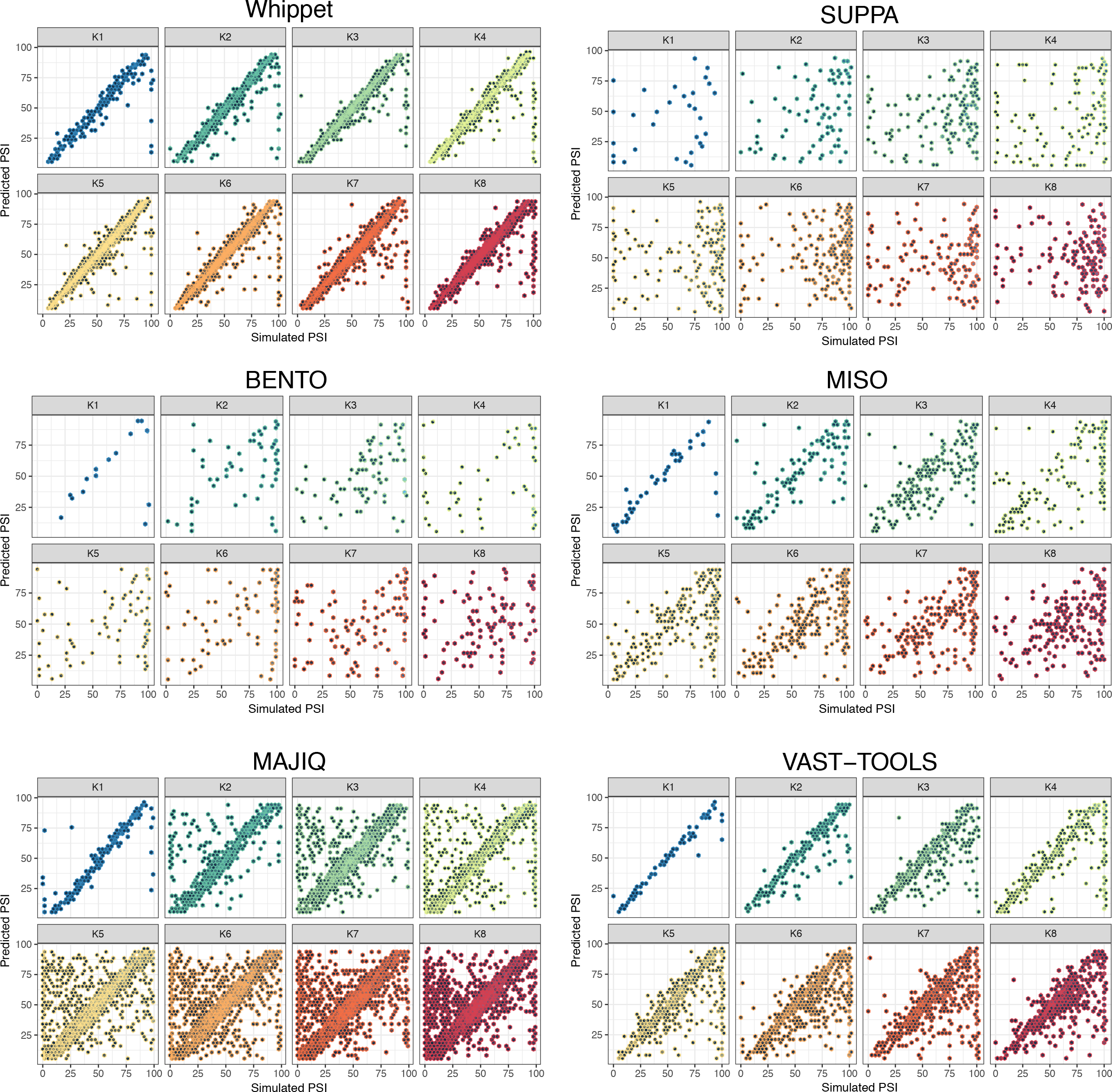
Scatter plots showing correlations between simulated PSI values and predicted PSI values from RNA-seq analysis by multiple published splicing algorithms at various levels of complexity (K*n*). (see Methods for details on data simulation).

**Supplementary Figure 6.**
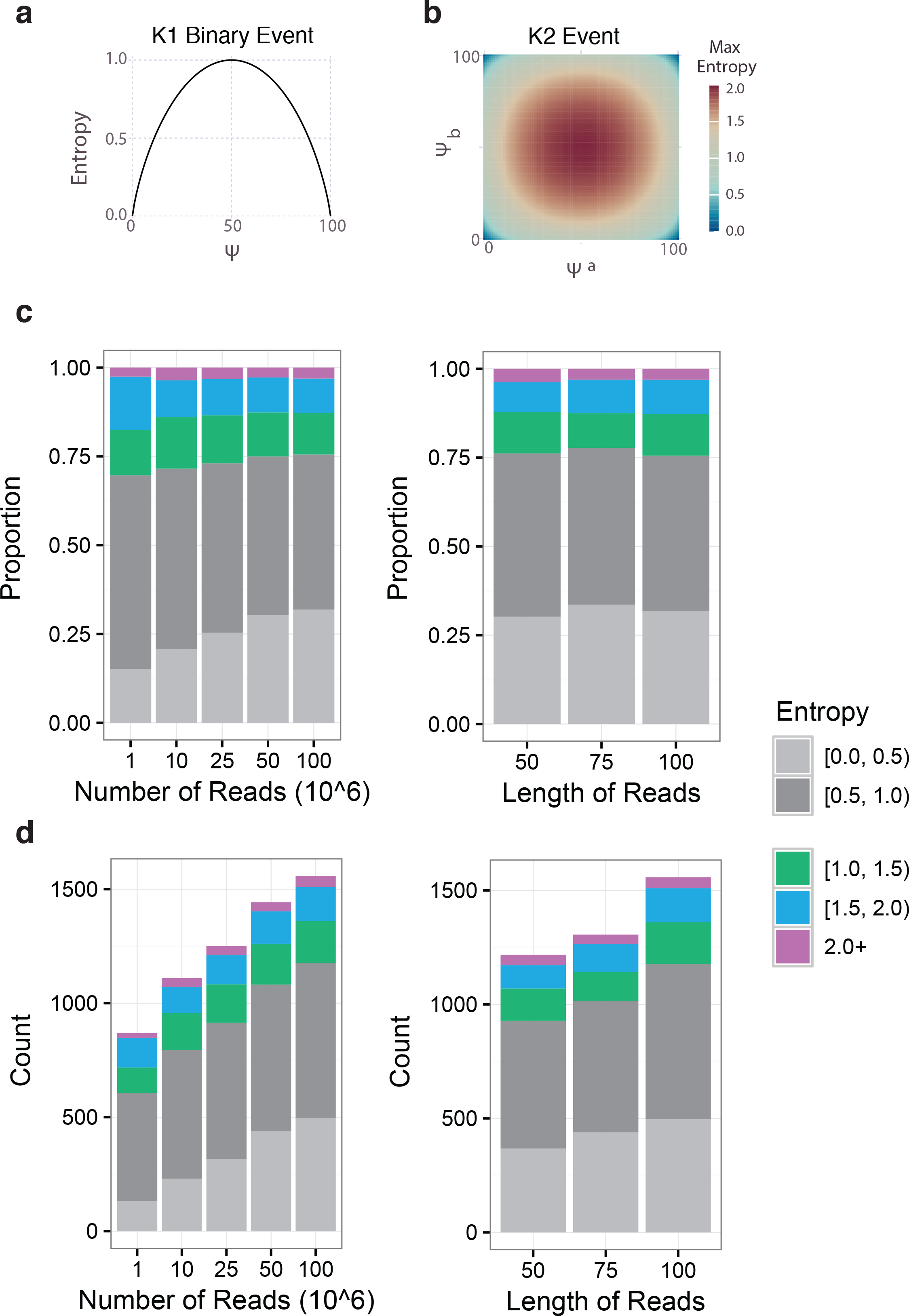
**(a)** Plot of entropy (*y-axis*) vs. percent-spliced-in (*x-axis*) for a simple binary (K1) AS event. **(b)** Plot of maximum entropy (*color-scale*) vs. percent-spliced-in (*x-axis,y-axis*) for a K2 event with two alternative exons *a* and *b* and two independent values for the percent-spliced-in of each exon, Ψ_a_ and Ψ_b_. **(c)** Proportion of AS events with entropy values in discrete ranges (*color-scale*) for the same simulated RNA-seq data set from sub-sampled read-depth in millions (*left*) and truncated read-lengths (*right*). **(d)** Total count of the number of AS events detected with the RNA-seq datasets from panel (c).

**Supplementary Figure 7.**
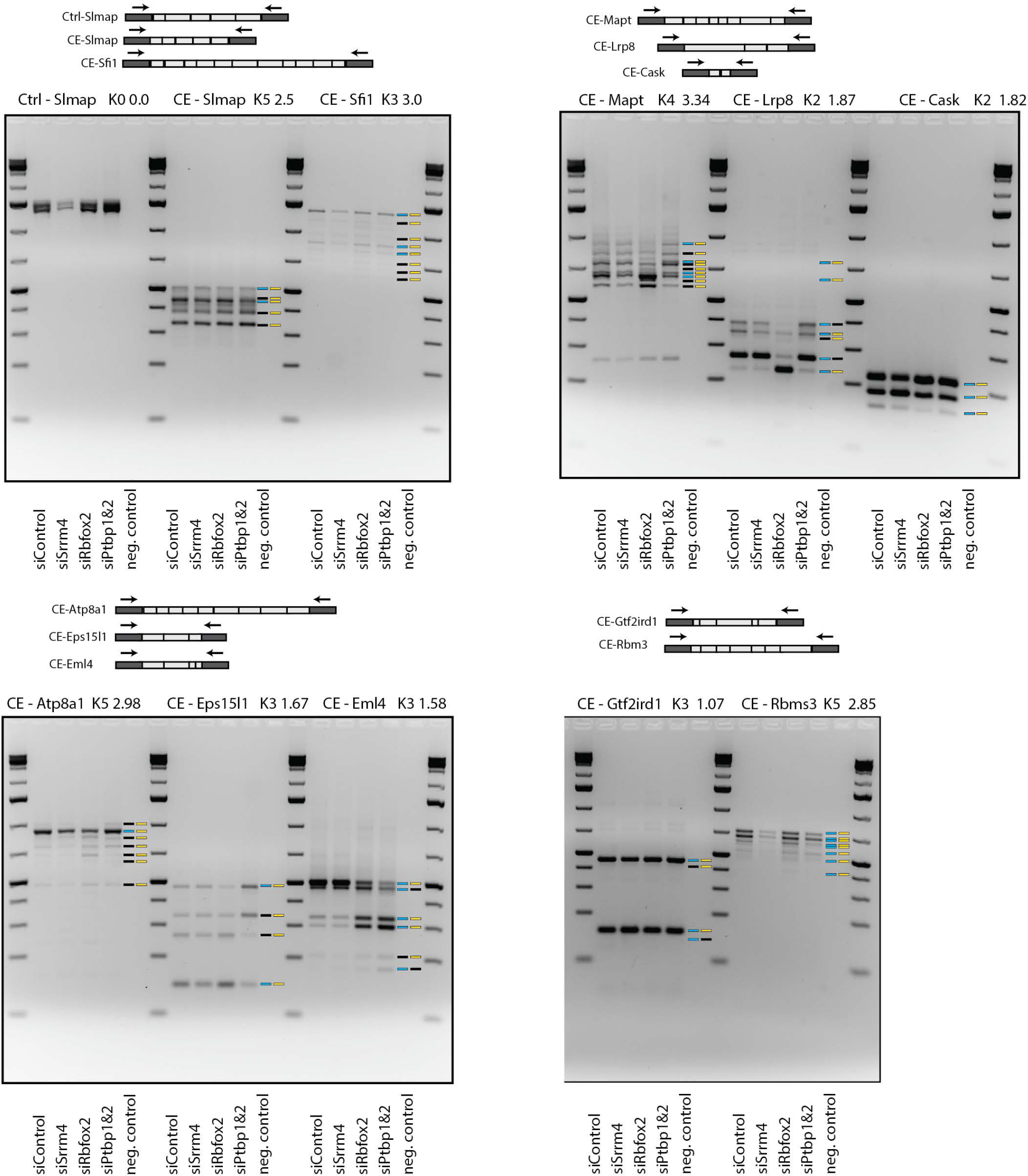
Extension of **Fig. 2c**, showing uncropped gels for events in the main figure as well as additional examples. RT-PCR analysis confirms the presence of complex splicing events in N2a cells at increasing levels of complexity matching Whippet predictions. Event type, gene name, complexity type and entropy score are shown above each events. Control SImap demonstrates that complexity is not just due to number of exons monitored. Boxes to right of gels display UCSC (*left*) and Whippet (*right*) predictions based on primer sequences (see Methods). Colored boxes represent correct predictions whereas black boxes suggest missed predictions. Diagrams below show exon structures of analyzed genes with approximate positions of RT-PCR primers indicated. Predicted constitutive and alternative exons are indicated in dark and light gray, respectively.

**Supplementary Figure 8.**
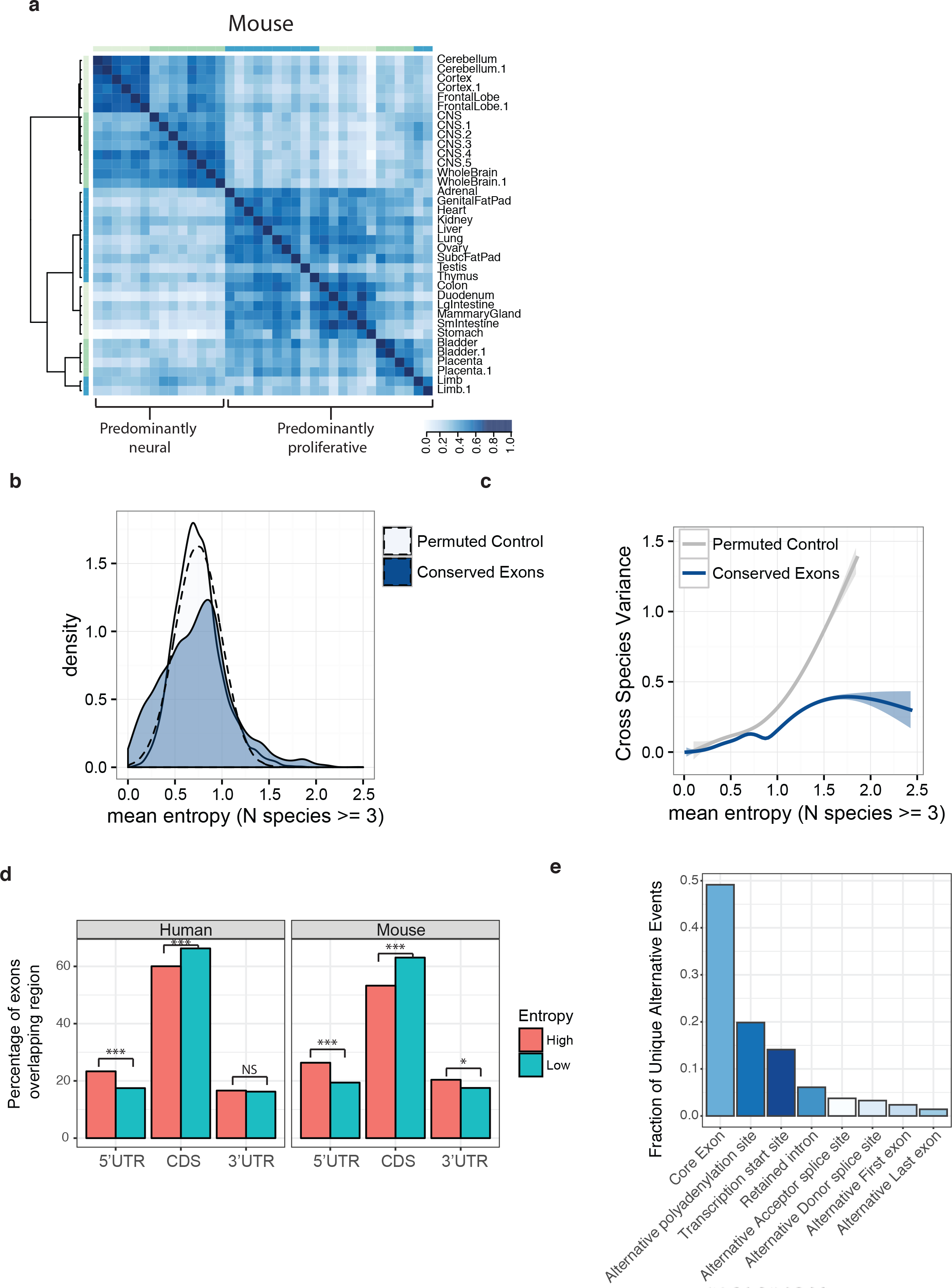
**(a)** Symmetrical heatmap of pairwise correlations of normalized splicing entropy scores across multiple mouse tissues. Heatmap showing affinity propagation clustering of pairwise similarities between splicing entropy scores. Colored bars surrounding heatmap indicate clusters defined by the dendrogram. Darker blue represents stronger correlation in splicing entropy between tissues types, whereas lighter blue indicates weak or no correlation. **(b)** Kernel density plots of mean entropy (*x-axis*) of AS events in the same tissue across at least three vertebrate species (human, chimp, gorilla, mouse, opossum, platypus, and chicken) for conserved exons (with liftOver) and permuted AS event labels for each species. Dashed density curve displays the best-fit Gaussian distribution to the permuted data. **(c)** Best fit ‘loess’ local smoothing for the variance of entropy (*y-axis*) vs. mean entropy (*x-axis*) for same AS events in (c). **(d)** Bar plots showing distribution of high entropy (>1.0) and low entropy (<1.0) events within 5’-UTR, CDS and 3’-UTR of transcripts across human and mouse tissues. **(e)** Bar plot showing fraction of unique alternative events identified by Whippet that can be detected from the multi-tissue human survey. All alternative events must be confidently identified with 0.05<PSI<0.95 except retained introns events when Percent Intron Retained (PIR) > 0.05. ***, p < 1.0 × 10−^05^ ; *, p<0.05; NS, not significant; CDS, coding sequence; UTR, untranslated region.

**Supplementary Figure 9.**
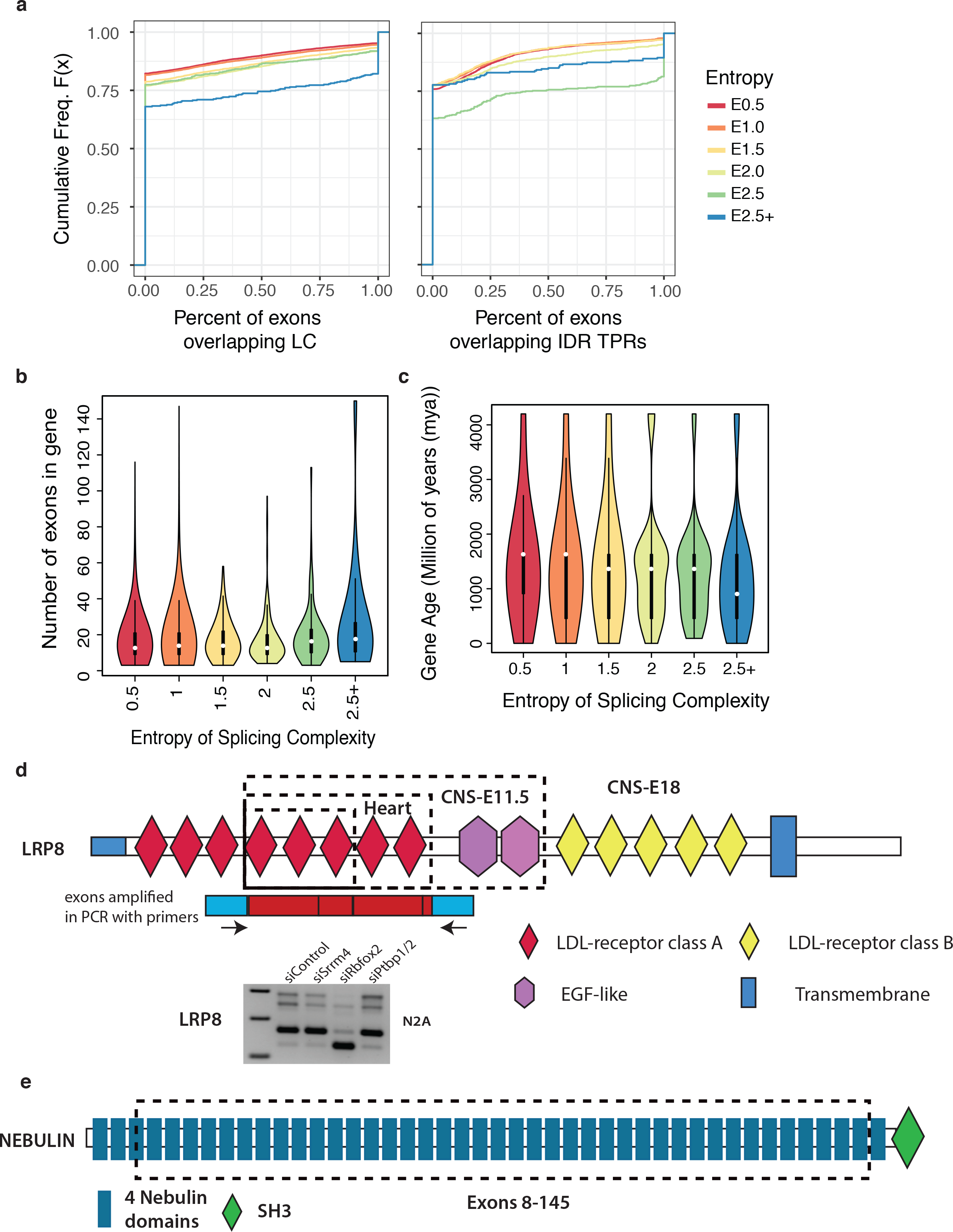
**(a)**Cumulative distribution plots showing: frequency of overlap of AS events (with different degrees of entropy) within (*left*) low complexity (LC) regions of proteins and (*right*) disordered tandem protein repeats (TPRs). See Fig. 2b legend for a description of cumulative distribution plots. (b) Violin plot showing the number of exons encoded by a gene body at different degrees of splicing entropy (maximum splicing entropy observed within gene body used to bin genes). See **Supplemental Fig. 9a** for color legend. See Fig. 2i for description of violin plots. **(c)**Violin plot showing that genes with higher complexity splicing events tend to be younger (or more recent gene duplication events). The violin plots include a marker for the median of the data, a box indicating the interquartile range and a visualization of the full distribution of the data. See **Supplemental Fig. 9a** for color legend. **(d)**Domain diagram for LRP8 (Low-density lipoprotein receptor-related protein 8) based on SMART annotation ^49^. Dotted boxes describe area of proteins undergoing high entropy splicing in different tissues types. Domain diagram below illustrates exons undergoing splicing within N2a cells and position of primers for RT-PCR validation below. **(e)**Domain diagram for Nebulin, based on SMART annotation ^49^, with dotted box indicating region undergoing highest entropy splicing.

**Supplementary Figure 10.**
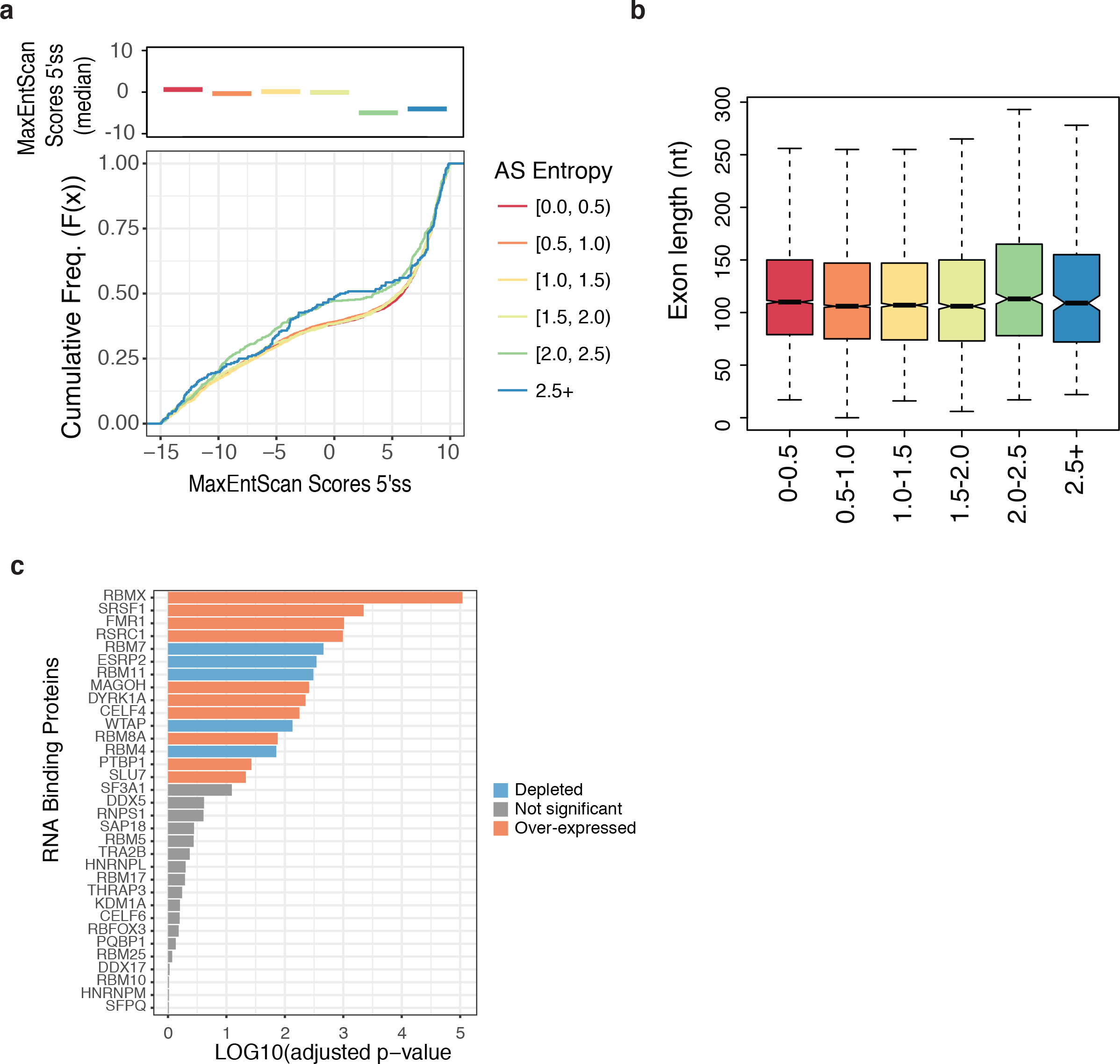
**(a)**Plots showing the cumulative distribution of 3’-splice site (3’ss) strength estimated using MaxEntScan^48^ and binned by different degrees of splicing entropy (*bottom*). The median 3’ss strengths for AS events with different degrees of splicing entropy are plotted as colored lines (*top*). See Fig. 2b for description of cumulative distribution plots. **(b)**Boxplot displaying lengths of introns surrounding exons binned by different degrees of splicing entropy. See **Supplemental Fig. 10a** for color legend. See Fig. 3d for description of box plots. **(c)**Full list of DESeq2 differential gene expression analysis50 between tumour samples and matched controls for selected RNA-binding proteins (GO:0000380). Genes with blue bars show reduced expression in cancer samples, red bars show increased expression in cancer samples, and gray bars show no significant difference between control and tumour samples

